# Coherent mapping of position and head direction across auditory and visual cortex

**DOI:** 10.1101/2021.07.30.452931

**Authors:** Paul EC Mertens, Pietro Marchesi, Matthijs Oude Lohuis, Quincy Krijger, Cyriel MA Pennartz, Carien S Lansink

**Affiliations:** Swammerdam Institute for Life Sciences, Center for Neuroscience, Faculty of Science, University of Amsterdam, Science Park 904, 1098 XH, Amsterdam, the Netherlands; Research Priority Program Brain and Cognition, University of Amsterdam, Science Park 904, 1098 XH, Amsterdam, the Netherlands

## Abstract

Neurons in primary visual cortex (V1) may not only signal current visual input but also relevant contextual information such as reward expectancy and the subject’s spatial position. Such location-specific representations need not be restricted to V1 but could participate in a coherent mapping throughout sensory cortices. Here we show that spiking activity in primary auditory cortex (A1) and lateral, secondary visual cortex (V2L) of freely moving rats coherently represents a location-specific mapping in a sensory detection task performed on a figure-8 maze. Single-unit activity of both areas showed extensive similarities in terms of spatial distribution, reliability and position coding. Importantly, reconstructions of subject position on the basis of spiking activity displayed decoding errors that were correlated between areas in magnitude and direction. In addition to position, we found that head direction, but not locomotor speed or head angular velocity, was an important determinant of activity in A1 and V2L. Finally, pairs of units within and across areas showed significant correlations in instantaneous variability of firing rates (noise correlations). These were dependent on the spatial tuning of cells as well as the spatial position of the animal. We conclude that sensory cortices participate in coherent, multimodal representations of the subject’s sensory-specific location. These may provide a common reference frame for distributed cortical sensory and motor processes and may support crossmodal predictive processing.

## Introduction

Early sensory cortical areas were long viewed to primarily function as collections of unisensory feature detectors (DiCarlo and Cox, 2007; Felleman and Van Essen, 1991; Hubel and Wiesel, 1962; Miller, 2016). More recently, single unit recordings in awake, behaving animals have shown responses in primary auditory (A1) and visual cortex (V1) to a wide variety of perceptual and behavioral factors, suggesting these areas have functions beyond unimodal sensory processing. In rodent V1, these include responses to reward and reward timing (Shuler and Bear, 2006), reward predictive stimuli (Goltstein et al., 2013), running speed (Ayaz et al., 2013; Niell and Stryker, 2010), head orienting movements (Guitchounts et al., 2020) and responses which are causal to visually cued action timing (Namboodiri et al., 2015). Additionally, a growing number of studies show that many V1 neurons display location-selective spiking activity (Ji and Wilson, 2007), coding the animal’s position along real (Haggerty and Ji, 2015) and virtual linear tracks (Fiser et al., 2016; Fournier et al., 2020; Pakan et al., 2018; Saleem et al., 2018). Various studies report spatial and temporal correlations in activity of V1 and hippocampal CA1, including correlated errors in position decoding (Fournier et al., 2020; Saleem et al., 2018), correlated trial-by-trial shifts in preferred spiking locations (Haggerty and Ji, 2015) and significant spike-phase coherence of V1 spiking and hippocampal theta oscillations (Fournier et al., 2020).

A similarly broad variety of single unit correlates is observed in A1, including activity selective for visual task-cues (Brosch et al., 2005), behavioral demands (Scheich et al., 2007), stimulus expectation (Jaramillo and Zador, 2011), reward (Scheich et al., 2007) and instrumental action (Niwa et al., 2012). Much remains currently unknown about the functional role and origins of such ‘extra-modal’ activity correlates, including whether they primarily contribute to local sensory processes or reflect crossmodal interactions in service of more general and modality-independent cortical functions. While spiking correlates to stimulus location are present in auditory cortex (Town et al., 2017), no activity selective for the spatial position of the subject has hitherto been reported for A1. This would be expected if the underlying mechanisms reflect general functions of cortical sensory processing, including the maintenance and updating of a coherent representation of space or of current and future sensory states across sensory domains (Fiser et al., 2016; Friston, 2005; Gavornik and Bear, 2014; Pennartz et al., 2019; Rao and Ballard, 1999).

To determine whether location-selective spiking activity exists outside of the visual cortical system and whether such activity provides a coherent representation across sensory modalities, we analyzed single-unit data recorded simultaneously from two anatomically connected, sensory cortical areas of freely moving rats: primary auditory cortex (A1) and lateral, secondary visual cortex (V2L). We show spatially localized firing patterns in large proportions of V2L and A1 single units that are reliable over time. Firing patterns in each area collectively tiled the entire behavioral track, so that every location was marked by activity of a subset of neurons. Reconstructions of the rat’s position afforded by the spiking activity of each area showed reconstruction errors that were correlated in magnitude and direction, thereby indicating that representations in A1 and V2L are coherent. Cross-areal coordination of location-specific representations was further indicated by strong noise-correlations in spiking activity of the vast majority of single-unit pairs which showed a clear pattern of dependence on both the spatial tuning of the units as well as animal position. Our freely moving paradigm allowed us to establish the contributions of position, head direction and their temporal derivatives to location-selective spiking activity in early sensory cortices dedicated to different modalities. Our results uncover striking similarities as well as quantitative differences in location-selective neural activity of A1 and V2L, suggesting that such activity supports common functions in coordinated mapping of sensory and contextual representations across different sensory modalities.

## Results

We investigated the responsiveness of neurons in lateral, secondary visual cortex (V2L) and primary auditory cortex (A1) to spatial location when rats were running on a rectangular, figure-8 shaped track (Fig.1A). On the track, rats performed an audio-visual discrimination task in which they earned reward by responding to the most salient stimulus out of two by running from the stimulus presentation site to the reward well on the track side corresponding to the location of that stimulus (Fig. 1 A-B). Our analyses were primarily based on the spatial components of the rats’ behavior, regardless of task performance. We refer to “location-selective activity” or a “location correlate” if the neuron’s spiking activity was reliably modulated by the rat’s body location over the course of a recording session. We emphasize that this definition includes not only responses to allocentric position, but also to specific conjunctions of locally available sensory cues and task-related information.

**Figure 1:**
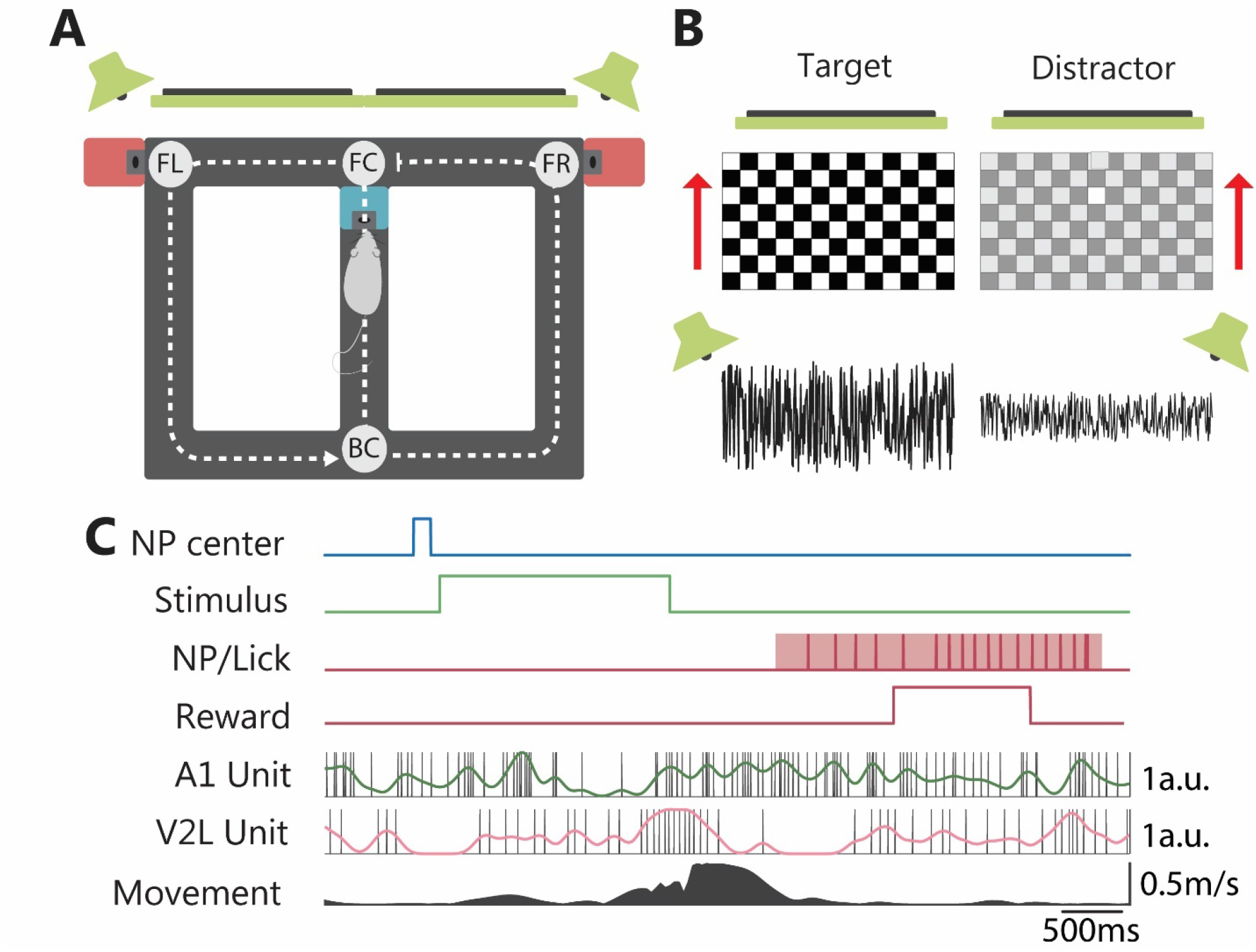
Behavioral apparatus and task. **(A)** Rats performed a discrimination task on an automatized figure-8 track, which was located in a dark, sound-attenuated laboratory room without other salient visual cues. At the front center T-junction (FC), rats responded to audio-visual stimuli presented from 2 screens and 2 speakers located in front of the track (green) by running to the track’s side corresponding to where the most salient stimulus was presented. Rats were rewarded for a correct response with sucrose solution at the ports to the sides of the front alley (pink squares). The dotted line indicates how the track was linearized for analysis. FL: front left, FR: front right, BC: back center. **(B)** Example of a set of stimuli. Stimuli were visual (moving checkerboard), auditory (filtered white noise) or audiovisual. In multisensory trials, target auditory and visual stimuli were always presented at the same side of the track. The rat had to respond to the most salient stimulus (highest contrast and/or volume) and discard the distractor stimulus (which was less salient). **(C)** Trial layout and example A1 and V2L spike trains during seven seconds of a leftward trial with a correct response. NP center: blue line indicates the timing of the nose poke (NP) in the central well to initiate stimulus onset. Stimulus: the time interval during which the stimulus was presented (2s; green line). NP/Lick: the time of the rat’s nose poke into the left reward well is indicated by the red shaded area and the individual licks by the vertical tick marks. A1 / V2L Units: Single unit spikes are indicated by the black vertical tick marks, whereas solid lines indicate Gaussian-smoothed spike trains. Firing rates are observed to fluctuate in relation to stimuli and locations across the maze. Movement: speed of the rat along the linear trajectory.

We made simultaneous recordings from both V2L and A1, with a total of 526 single units from V2L and 603 units from A1 across 17 recording sessions from 3 rats. Units in both A1 and V2L showed firing patterns with one or more peaks in firing rate on various locations of the track (Fig. 1C, 2A-B). For all further analyses, we selected the units with sufficient firing on the track, i.e. peak firing rate above 2 Hz in the spatial map, which amounted to 400 units from V2L and 413 from A1. The locations of the tetrode endpoints were verified with histology (Supplementary Fig. 1). The endpoints of 20 tetrodes were located in A1, while an additional two endpoints targeted at A1 were located in the adjacent dorsal secondary auditory cortex. Endpoints for 19 tetrodes targeted at V2L were located in that area, while one was located in the adjacent dorsal posterior parietal cortex.

**Figure 2:**
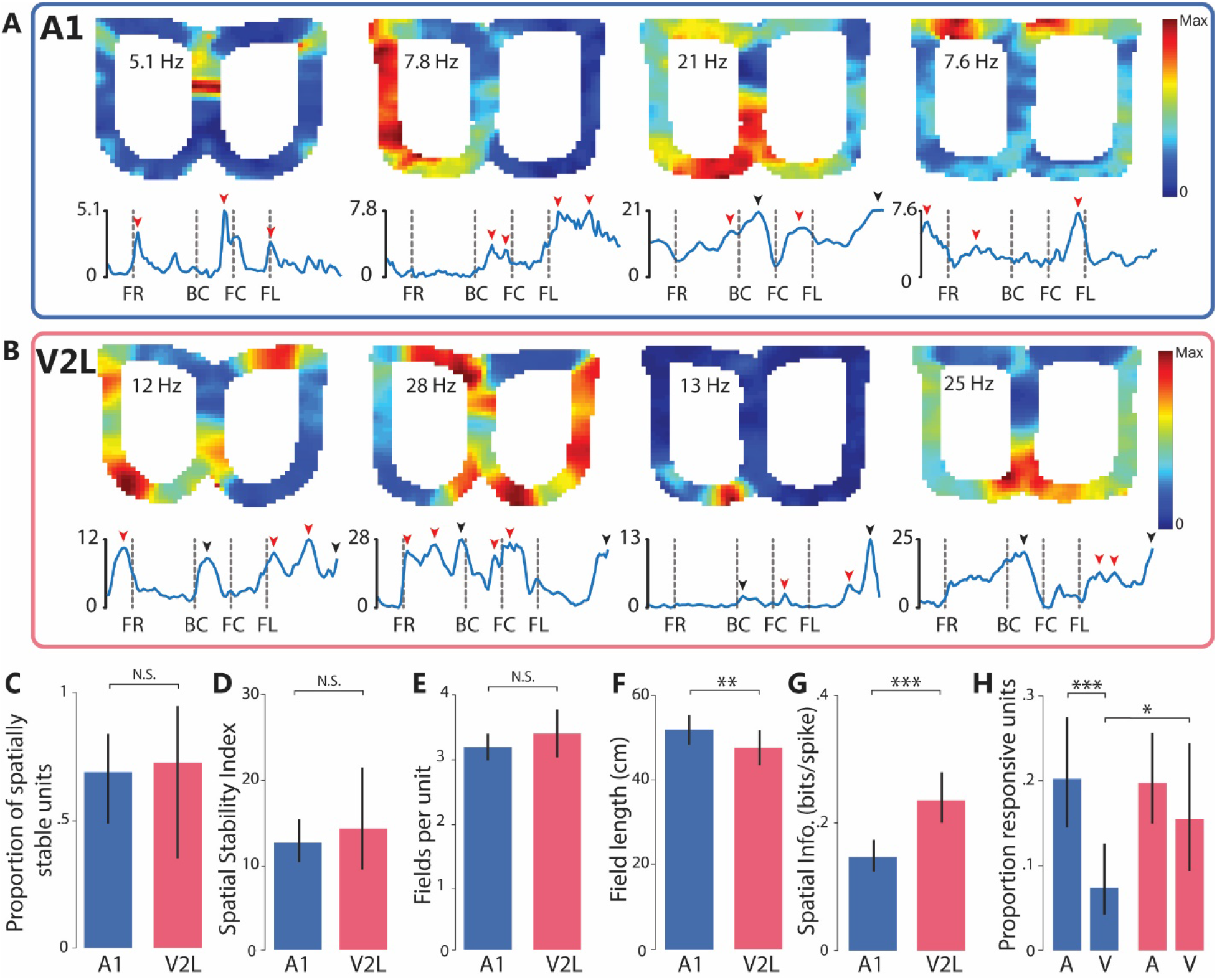
Spatially localized firing patterns of A1 and V2L units. **(A)** *Top panels.* Rate maps indicating the spatial firing-rate distribution of four example A1 units. The firing rate of the units is color-coded and the peak rates are indicated in the left wing of the map. *Bottom panels.* Firing rate distributions along the linearized version of the track corresponding to the rate maps shown in the top panels. Red arrow heads indicate individual firing field peaks. Black arrow heads indicate a single firing field peak that spans across an edge of the linearized track. Abbreviations below linear rate maps refer to landmark locations (fig. 1A). **(B)** as (A) but for four example V2L units. **(C)** Estimate of the proportion of spatially stable units in the A1 and V2L populations. Black vertical lines throughout (C-H) indicate 95% confidence intervals (N.S.: p > 0.05, F-test). **(D)** Spatial stability index of spatially stable units. **(E)** Number of firing fields per spatially stable unit **(F)** Field length per unit (**, p < 0.01). **(G)** Spatial information of spatially stable units (***, p < 0.001). **(H)** Proportions of stimulus responsive neurons in A1 (blue bars) and V2L (red) to auditory and visual stimuli (*, p < 0.05).

### The majority of A1 and V2L single units displays localized spiking activity

First, we assessed whether neurons in A1 and V2L displayed spatially localized firing patterns. Visual inspection of rate maps of the spatial firing distribution of individual neurons indicated that some units in A1 and V2L showed increased spiking activity in a single, concentrated location on the track, whereas other units displayed modulations of spiking activity at multiple areas or across a larger area of the track (Fig. 2A-B *top panels*). An important constraint for a unit to reliably code location is that the firing activity at that location is consistently, rather than incidentally, present across individual traversals through that location. Therefore, we assessed the reliability of each unit by computing pairwise correlations between all single-trial rate maps of that unit and comparing the observed mean pairwise correlation with a shuffled distribution. A unit was considered spatially stable if its observed mean correlation was larger than 95% of the shuffled distribution. The proportions of spatially stable units were very similar for A1 and V2L and comprised on average about 70% of the total population (Fig. 2C; A1: 0.69 (CI: 0.49, 0.84), V2L: 0.72 (CI: 0.31, 0.94), F(1, 32) = 0.072, p = 0.79, ANOVA). The degree of spatial stability of a unit can be quantified using a spatial stability index (SSI), defined as the number of standard deviations which the observed mean correlation was removed from the shuffled distribution (Fig. 2D). The SSI of spatially stable units was similar between A1 and V2L (Fig. 2D; A1: 12.7 a.u. (CI: 8.39, 15.44), V2L: 14.3 a.u. (CI: 10.4, 24.4), F(1, 1194) = 0.44, p = 0.50, ANOVA). In summary, large fractions of A1 and V2L units display spatially stable activity patterns with high reliability.

Individual firing fields were then identified for spatially stable units as localized increases in mean spiking activity of a unit on the linearized representation of the track (Fig. 1A, Fig. 2A-B). Whereas neurons in A1 and V2L exhibited a similar number of around 3 firing fields per unit (Fig. 2E; A1: 3.17 (CI: 2.97, 3.39), V2L: 3.40 (CI: 3.20, 3.61), F(1, 588) = 2.22, p = 0.14, ANOVA), the average length of firing fields was significantly smaller in V2L than in A1 (Fig. 2F; A1: 50.5 cm (CI: 47.2, 54.1), V2L; 46.3 cm (CI: 43.5, 49.4), F(1, 1939) = 9.0, p = 0.003, ANOVA), indicating that the spatial granularity of localized spiking activity is modestly finer in V2L than A1.

The extent to which units show spatially localized firing can be expressed as the information about the rat’s position that is conveyed by a single spike, which is quantified as spatial information (Skaggs et al., 1992). In line with the smaller firing fields in V2L, the spatial information was significantly higher for V2L units (Fig. 2G; A1: 0.15 bits/spike (CI: 0.12, 0.17), V2L: 0.24 bits/spike (CI: 0.20, 0.28), F(1,556) = 16.5, p = 5⁎10^−5^, ANOVA).

### Both A1 and V2L neurons respond to discrete auditory and visual stimuli

Besides showing location-selective firing, subsets of units in both A1 and V2L responded with significant firing-rate changes to the auditory and visual stimuli, presented as individual (unisensory) task cues (Fig. 2H; stimulus conditions were pooled for each modality; see Methods). As expected, a larger proportion of A1 neurons responded to auditory than to visual stimuli in all rats (A1 auditory: 0.20 (CI: 0.15, 0.28), A1 visual: 0.07 (CI: 0.04, 0.13), F(1, 30) = 10.34, p = 0.003, ANOVA). Additionally, responsiveness to visual stimuli was more common in V2L than A1 (V2L visual: 0.15 (CI: 0.09, 0.24), F(1,30) = 4.60, p = 0.0402, ANOVA). Surprisingly, however, a comparable proportion of V2L neurons responded to both visual and auditory (stimuli. (V2L auditory: 0.20, (CI: 0.15, 0.26), F(1, 30) = 0.93, p = 0.34, ANOVA) and auditory responses were equally common in both areas (F(1, 30) = 0.013, p = 0.91, ANOVA). Although A1 and V2L responsiveness to discrete stimuli is not the focus of our current analyses, these results not only indicate that responses to stimuli in more than one modality are common in both A1 and especially V2L, but also underscore the existence of substantial heterogeneity in cortical sensory selectivity. Finally, it should be noted that stable, localized activity was much more abundant in both areas than responses to sensory stimuli (A1: p < 10^−15^; V2L: p < 10^−13^, binomial tests on pooled data).

### Spatial firing field distributions are highly correlated across A1 and V2L

The reliable coding of specific locations by individual units is necessary, but not sufficient for building a representation of an environment or the sequence of sensory states an animal experiences when navigating across the track. Another prerequisite for either type of representation would be that the spatial distribution of firing fields in each area covers the entire series of locations traversed by the animal. Figure 3A and B show the linear rate maps for all spatially stable units ordered by peak location for A1 and V2L, and reveal that firing-field peaks occur at every location along the track. We further analyzed the density of firing fields tiling all locations by computing the proportion of firing fields which include a specific spatial bin on the linearized track. The distributions of firing field densities confirm that the entire track was covered by A1 and V2L firing fields (Fig. 3A-B, bottom).

**Figure 3:**
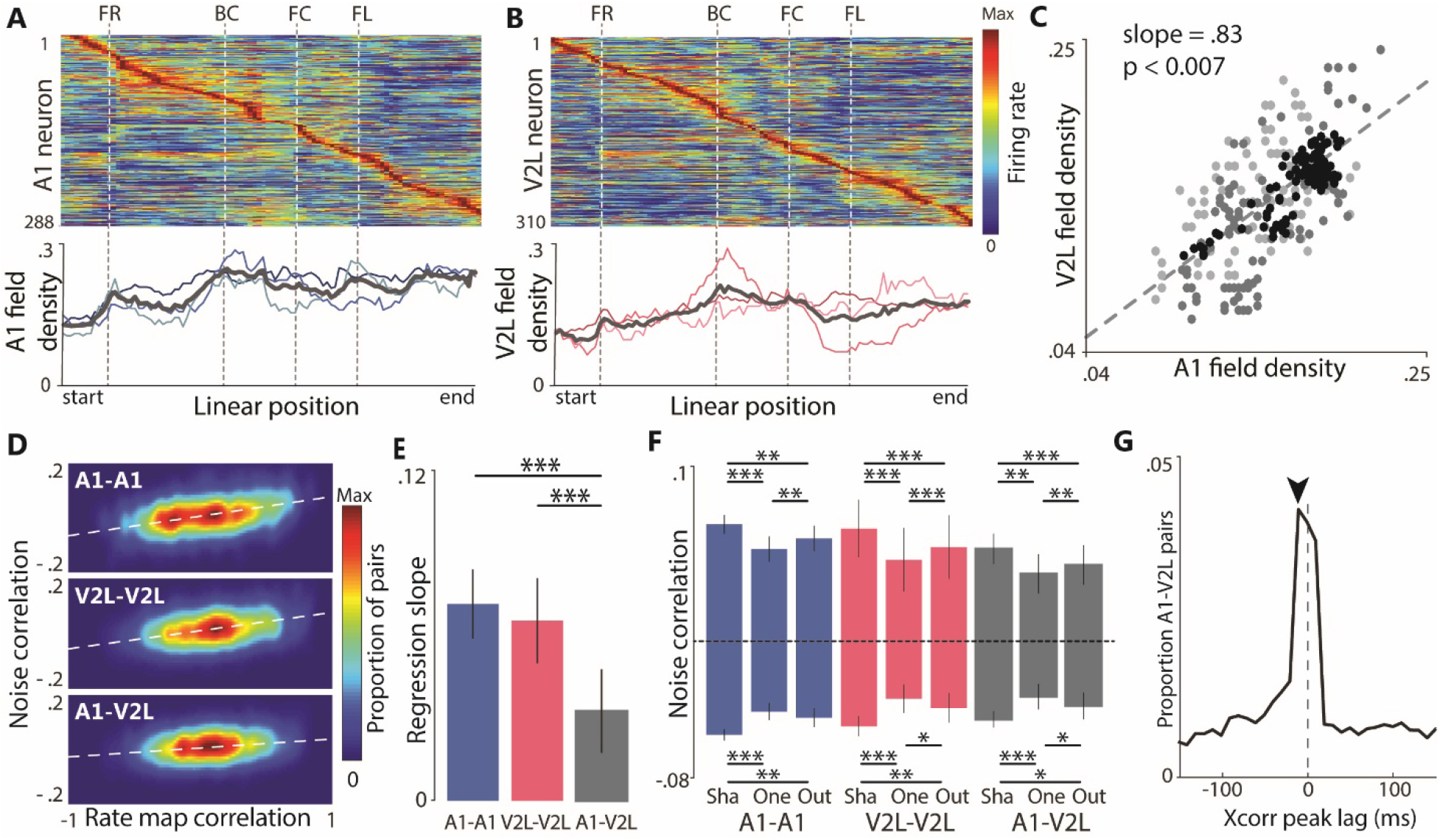
Spatial and temporal firing relations between A1 and V2L neurons. **(A)** *Top panel.* Joint linear rate map including all individual, spatially stable units recorded from A1. Firing rate is color-coded and the individual rate maps are sorted by peak location. *Bottom panel.* The firing field density across the spatial bins on the track are shown for the individual rats (thin, colored lines) and the mean across rats (thick, grey lines). The firing field density expresses for each spatial bin the proportion of individual firing fields that include that bin. **(B)** as (A) but for V2L. **(C)** Correlation between firing field densities of A1 and V2L across spatial bins. Dots correspond to individual spatial bins; different shades of grey correspond to individual rats. The dashed line indicates the linear regression of mean field densities across rats according to the linear mixed effects model. The value of *p* indicates the significance of the regression line. **(D)** Heat maps show the proportions of cell pairs averaged across rats as a function of observed rate map correlations and noise correlations for A1-A1 pairs (top), V2L-V2L pairs (middle) and A1-V2L pairs (bottom). White, dashed lines indicate regression lines from a linear mixed effects model, indicating the significant, positive relationship between rate map correlations and noise correlations. **(E)** Bars indicate the slope of the regression of noise correlations on rate map correlations (white, dashed lines in **D**) for the different single-unit pair types, with error bars indicating 95% confidence bounds versus 0. Asterisks indicate a significant difference between groups (***: p < 0.001). **(F)** Mean pairwise noise correlations for the within and between-area pairs and field configuration presented for significantly positively correlated pairs and significantly negatively correlated pairs separately. *Sha*: locations shared between firing fields of both units; *One*: locations exclusively appearing in the firing field of one of the two units, but not the other; *Out*: locations outside of the firing fields of either unit. Error bars are 95% confidence bounds versus 0. Asterisks indicate a significant difference between groups (***: p < 0.001, **: p < 0.01, *: p<0.05). **(G)** Proportions of A1-V2L unit pairs with observed temporal cross-correlation peak lags, averaged across rats. Black arrow indicates the lag with the highest average proportion of pairs. Negative lags indicate that activity of the A1 unit precedes that of the V2L unit.

The joint rate maps and field density plots indicate similar firing-field distributions in A1 and V2L. Indeed, the field densities across spatial bins of the track were highly correlated between the areas (Fig. 3C; regression slope: 0.83 (CI: 0.57, 1.09), F(1, 39.5) = 3.2p = 0.007, ANOVA). This strong correlation, however, may also be due to similar responses of A1 and V2L neurons to behavioral covariates such as locomotion speed, acceleration and head direction (θ_head_). To correct for the potential influence of these covariates on field densities, we performed linear regression of spiking activity on running speed, acceleration, head direction and the angular change in head direction (Δθ_head_) and repeated the detection of firing fields and the field density correlations on the residuals. Likewise, when using the model’s residuals, field densities between A1 and V2L were highly correlated (regression slope: 0.71 (CI:0.36, 1.06), F(1, 3.0) = 16.1, p = 0.028, ANOVA). It is therefore unlikely that locomotion and head direction can fully explain the firing-field densities of A1 and V2L neurons on the track or their shared spatial distribution. These analyses, however, did not take into account possible nonlinear relationships between location and behavioral covariates.

### Temporal firing rate fluctuations are correlated between A1 and V2L

Neurons with similar functional properties, such as location-selective tuning in hippocampal and V1 neurons, were shown to exhibit significant pairwise correlations of the instantaneous variability in spiking activity, often referred to as ‘noise correlations’ (Haggerty and Ji, 2015). As a first indication of a correlative interaction between spatially stable neurons within and across A1 and V2L, we tested how closely the precise firing rate between two units of a pair varied each time the rat ran through the associated firing fields. For each pair of units (A1-A1, V2L-V2L, A1-V2L) the Pearson correlation was computed between their instantaneous variability in spiking activity after subtraction of the mean activity in the associated spatial bin (i.e. noise correlation), as well as the Pearson correlation between the rate maps. Noise correlations were significantly different from 0 for very large proportions of neuronal pairs of each composition (fraction of total: A1-A1: 0.98, V2L-V2L: 0.98, A1-V2L: 0.97; permutation test, p<0.05). Almost half of the significantly correlated pairs showed negative noise correlations (A1-A1: 0.46, V2L-V2L: 0.47, A1-V2L: 0.45). Proceeding only with pairs showing significant noise correlations, we tested whether rate map similarity predicted noise correlations using a linear mixed model and correcting for sampling bias. Rate-map correlations had a significant, positive effect on noise correlations for all pair types (Fig. 3D-E; regression slopes and 95% confidence bounds for A1-A1: 0.072 (0.060, 0.085), F(1, 4.8) = 123.9, p =3.9 ⁎ 10^−4^, V2L-V2L: 0.066 (0.051, 0.082), F(1, 4.17) = 110.4, p = 7.4 ⁎ 10^−4^, A1-V2L: 0.033 (0.018, 0.048), F(1, 3.67) = 29.3, p = 0.007, ANOVA). The effect of rate map similarity was smaller for A1-V2L pairs than for the other pair types (A1-V2L slope vs. A1-A1 slope: F(1, 9386) = 95.9, p < 10^−15^, A1-V2L slope vs. V2L-V2L slope: F(1, 10174) = 83.2, p< 10^−15^, ANOVA), which is in line with previous observations on noise correlation strength decreasing with anatomical distance observed in visual cortex of anesthetized cats, mice and macaques (Goltstein et al., 2015; Rosenbaum et al., 2017; Schulz et al., 2015; Smith and Kohn, 2008). Nonetheless, the V2L-A1 noise correlates and their predictability from rate-map correlations suggests cross-areal coordination of neural activity.

### Noise correlations within and between areas show location-dependent structure

Correlated firing rate fluctuations can affect the coding of information in populations of neurons (Ecker et al., 2010; Hansen et al., 2012; Montijn et al., 2016; Zohary et al., 1994). The strength of correlated variability in the visual system was found to depend on both the tuning of neurons and the presented stimuli, with similarly tuned neurons displaying stronger correlated variability specifically for their preferred stimuli and reduced correlations for orthogonal stimuli, thereby ameliorating the potentially detrimental effect of correlated variability on population coding (Averbeck et al., 2006; Franke et al., 2016; Lin et al., 2015; Montijn et al., 2016). If the function of location-selective activity in A1 and V2L is to transmit information related to the animal’s position to downstream areas, correlated variability between neurons in both areas may show similar dependence on spatial tuning and location types. To study this, for each significantly correlated pair we divided all spatial bins of the track into three categories: bins that occur inside firing fields of both units of the pair (‘shared field’), bins that exclusively occur in a firing field of one of the two units (‘one-field’), and bins that are outside of firing fields of both units (‘out of field’). The effect of firing-field overlap on noise correlations for each area was examined using a linear mixed model. Separate models were constructed for positive and negative noise correlations, since the mean noise correlations of individual groups tended to 0, while a model based on absolute noise correlations provided a poorer fit to the data.

The magnitude of noise correlations followed a consistent pattern across cell pair types with respect to the type of bin, with shared field bins showing stronger correlations than one-field bins (figure 3F, positive correlations, shared field bins mean = 0.061, one-field bins mean = 0.046, F(1, 3.2) = 139.5, p = 4.7 ⁎ 10^−4^, negative correlations, shared field bins mean = −0.049, one-field bins mean = −0.035, F(1, 8.9) =233.9, p = 3.5 ⁎ 10^−8^, ANOVA). Additionally, shared field bins showed stronger correlations than out of field bins (figure 3F, positive correlations: out of field bins mean = 0.052, F(1, 2.9) = 43.3, p = 0.008, negative correlations: out of field bins mean = −0.040, F(1, 2.7) = 41.9, p = 0.005, ANOVA). Finally, out of field bins showed stronger correlations than one-field bins (figure 3F, positive correlations: F(1, 29.3) = 28.7, p = 3.0 ⁎ 10^−6^, negative correlations: F(1, 2.7) = 14.8, p = 0.04, ANOVA). These findings underscore the general result that noise correlations are highest in shared field bins and lowest in one-field bins, with noise correlations in out of field bins being intermediate. The only contrast not obeying this general effect is the comparison between negative correlations of one-field bins and out of field bins for A1-A1 pairs, which did not reach significance (p = 0.10).

The strength of noise correlations was found to depend modestly on the area-based type of cell pair, with A1-A1 pairs showing stronger noise correlations than A1-V2L pairs (figure 3F, positive correlations: A1-A1 mean = 0.059, A1-V2L mean = 0.045, F(1, 3.1) = 22.6, p = 0.048, negative correlations: A1-A1 mean =-0.046, A1-V2L mean = −0.038, F(1, 5.2) = 55.78, p = 0.002, ANOVA). Mean noise correlations for V2L-V2L cell pairs were intermediate to A1-A1 pairs and A1-V2L pairs, for both positive (V2L-V2L mean = 0.055) and negative correlations (V2L-V2L mean = −0.040), and differences between V2L-V2L pairs and the other cell pairs were not significant (all p > 0.05). There was no significant interaction between the type of bin and the composition of the neuronal pairs (positive correlations: F(1,14587) = 1.53, p = 0.19, negative correlations: F(1, 8312) = 1.59, p = 0.18, ANOVA). We found qualitatively similar results when we defined location type not in terms of falling inside or outside of firing fields of the two units of a pair, but in terms of falling inside the 30% of bins with highest or lowest spiking activity of the two units of a pair. To summarize, A1 and V2L populations do not only display similar distributions of firing fields across the maze, but pairs of A1 and V2L neurons also show fine-grained firing rate co-fluctuations which depend on both the similarity of location-selectivity of the units in a pair and the actual position occupied by the rat. Despite area-dependent differences in noise correlations, the similarity of A1-V2L noise correlation patterns to within-area co-fluctuations is rather striking.

### A1 activation precedes V2L activation in time

The fine-grained co-fluctuations between A1 and V2L in the spatial domain raise the question how these interactions are more precisely organized in time. The temporal firing relation between A1-V2L neuronal pairs was determined by computing the cross-correlograms over periods in which the rat was running on the track (running speed > 0.06 m/s). To correct for the impact that the forced unidirectional running may induce in estimating the temporal firing relation, cross-correlations were computed on the z-scored spiking activity after subtracting the mean activity in the associated spatial bin. Figure 3G shows the proportions of cross-correlation peaks of all A1-V2L pairs at different time lags averaged over rats. The narrow peak centered at −10 ms indicates that activation of A1 neurons mostly precedes V2L neuronal activity closely in time.

### Spiking activity carries information about position and head direction

We next used information-theoretic measures to quantify the influence of navigation parameters, i.e. position, running speed, head direction (θ_head_) and changes in head direction (Δθ_head_), on the spiking activity of recorded units. Mutual information (MI) quantifies the reduction in uncertainty obtained about spiking activity after observing these factors and captures both linear and nonlinear relationships (Lizier, 2014; Olcese et al., 2016). Of these individual factors, position carried the most information about spiking activity of both A1 and V2L units, followed by head direction (Fig. 4A; MI of position versus θ_head_: A1: p = 1.1 ⁎ 10^−15^; V2L: p =3.0 ⁎ 10^−15^, Wilcoxon signed-rank test). Running speed and changes in head direction carried relatively little information about spiking activity in both areas. For all individual factors mean MI was higher in V2L than in A1 (p = 2.0 ⁎ 10^−6^; running speed: p = 4.7 ⁎ 10^−4^, θ_head_: p =1.4 ⁎ 10^−7^, Δθ_head_: p = 1.6 ⁎ 10^−5^, Mann-Whitney’s U test). When a particular θ_head_ is predominantly encountered at a given position, mutual information cannot distinguish between the possibilities of a neuron encoding either that position, head direction or a combination of the two. To determine the amount of information carried by spikes about speed, θ_head_ and Δθ_head_, that cannot be explained by (nonlinear) correlations with position, we computed the debiased conditional mutual information (cMI) between each of the three factors conditional on position (Fig. 4B; (Bos et al., 2019; Lizier, 2014)). Averaged across all spatially stable units, the amount of information that spike trains carried about running speed and changes in head direction beyond position (i.e. mean cMI(spikes, speed | position) and mean cMI(spikes, Δθ_head_ | position)) was negative for both A1 and V2L, confirming that running speed and changes in head direction contributed little information about spiking activity on top of position (fig. 4B). In contrast, the average cMI for head direction, cMI(spikes, θ_head_ | position), was significantly larger than zero for both A1 and V2L (fig. 4B, A1: p < 10^−15^; V2L: p < 10^−15^; Wilcoxon’s signed-rank test), indicating that spiking activity of both A1 and V2L units coded information about head direction in addition to information about position.

**Figure 4:**
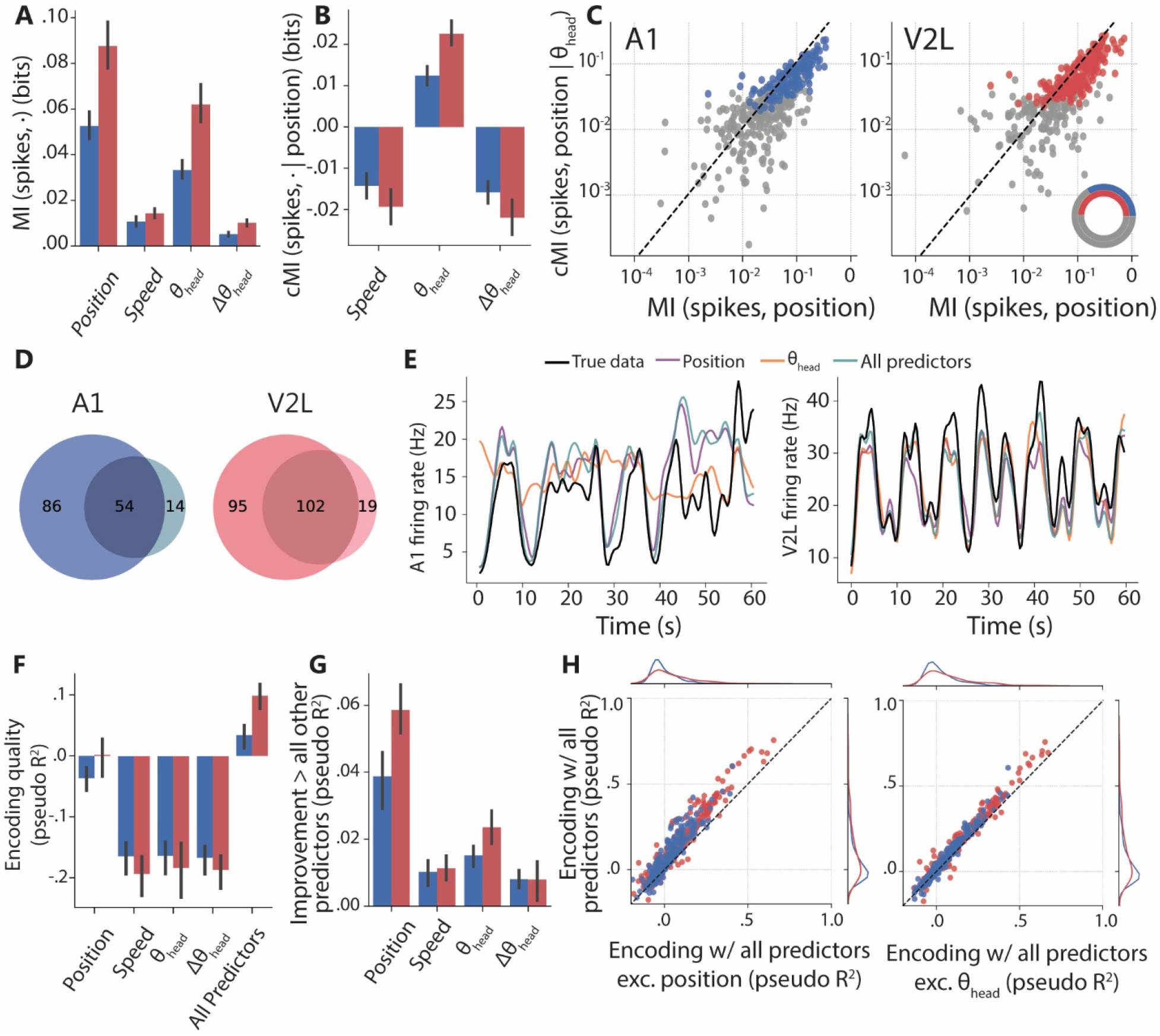
Encoding: predicting single-unit spiking activity from behavioral factors. **(A)** Average mutual information (MI) between spiking activity of A1 (blue) and V2L (red) single neurons and the behavioral factors position, running speed, head direction (θ_head_), and head direction change (Δθ_head_). Error bars represent 95% bootstrapped confidence bounds. **(B)** Debiased, conditional mutual information (cMI) between spiking activity and behavioral factors speed, head direction and changes in head direction, conditional on position. Error bars as in A. **(C)** Relationship between the MI of spikes and position and the cMI between spikes and position conditional on head direction for all spatially stable single units for A1 (left) and V2L (right). Blue/red points mark units with significant cMI about position conditional on θ_head_, indicating that these units carry significant information on position that cannot be explained by θ_head_. Inset shows in color the number of units per area showing significant cMI about position conditional on θ_head_ as fraction of the total number of units (grey). **(D)** Venn diagrams showing for each area the number of neurons transmitting significant cMI(spikes, position | θ_head_) (dark color) and cMI(spikes, θ_head_ | position) (light color). Overlapping region indicates neurons which transmit significant information on both position and θ_head_, neither of which can be explained entirely by the other factor. **(E)** A random forest encoder was used to predict spiking behavior on the basis of position, running speed, θ_head_ and Δθ_head_. Figure shows 60 s of firing rate of an example A1 unit (black line; left panel) and V2L unit (black line; right panel) and the predicted firing rate based on the models including the different behavioral parameters (colored lines). **(F)** Mean encoding quality across all A1 (blue) and V2L (red) units using single behavioral factors as predictor and using all predictors. Error bars as in A. **(G)** Mean improvement in encoding quality across all single units following the addition of the indicated behavioral factor to a model already containing all other factors. Error bars are 95% confidence bounds. **(H)** The relationship between encoding quality of individual single units when all predictors are considered and the encoding quality when all predictors except head direction (left) or linear position (right) are considered. Points above the diagonal belong to units with improved encoding due to the inclusion of linear position/ head direction which cannot be attributed to any other included factor. Blue: A1 units, red: V2L units. Diagrams to the right and top of the main scatterplot show the empirical distributions of the data depicted in the scatterplots projected onto a single dimension.

To study the prevalence of position coding in populations of single units, we calculated cMI between spikes and position conditional on θ_head_, thereby excluding any contributions from θ_head_ (Fig. 4C). Significant cMI (spikes, position | θ_head_) was found in 140 out of 413 A1 units (33%) and 197 out of 400 V2L units (49%), indicating that spiking activity of large proportions of single units in both areas contains information on spatial position on top of any information on θ_head_, with V2L showing a significantly larger proportion (z = 4.64, p = 1.7 ⁎ 10^−6^, binomial test; Fig. 4D). Smaller, but substantial subsets of neurons in each area showed significant cMI (spikes, θ_head_ | position), indicating that the spike trains of these neurons carry information about head direction even after correcting for position (fig. 4D, A1: 16.5% and V2L: 31.2%), with also here V2L showing a significantly larger proportion of neurons (z = 4.9, p = 4.2 ⁎ 10^−7^, binomial test). Among all units encoding position and/or head direction, V2L units were also significantly more likely to provide information on both factors (fig. 4D, A1: 35.1%, V2L: 47.2%, z = 2.33, p = 0.010, binomial test). In summary, this analysis confirmed that substantial fractions of A1 and particularly V2L neurons continued to show location- and head direction selectivity after correcting for the influence of the other behavioral covariate.

### Position is the strongest predictor of A1 and V2 single unit activity, followed by head direction

Next we used a random forest encoding algorithm (Benjamin et al., 2018) to determine how well spiking activity could be predicted by each of the behavioral factors. This provided the opportunity to extend the analysis of dependencies between behavioral factors and spiking activity beyond two behavioral factors. Encoding of firing rates of two units from A1 and V2L is exemplified in Figure 4E. For both areas, encoding quality was best for a model incorporating all predictors (Fig. 4F; all predictors vs. only position for A1 and V2L combined, p < 10^−15^, Wilcoxon’s signed-rank test), whereas of the individual factors position provided the best performance (position vs. each other individual predictor for A1 and V2L, all p < 10^−15^, Wilcoxon’s signed-rank tests). The large effect of position on encoding performance could not be explained by confounds arising from the other behavioral factors, since adding position to a model already utilizing all other factors led to substantial improvements in encoding performance for both A1 and V2L (fig. 4G; position vs. each other individual factor, all p < 10^−15^, A1 and V2L combined, Wilcoxon’s signed-rank tests). Comparing the contributions of position and θ_head_ separately for individual units confirmed that among the behavioral factors considered, position was dominant in explaining spiking activity in both areas, while θ_head_ provided the second strongest contribution (fig. 4G). This is also illustrated in Figure 4H, which shows for each neuron the improvement in encoding when adding θ_head_ or position to models containing all other behavioral factors. For the vast majority of units in both areas, adding either factor improved encoding quality, but this difference was larger for position than for θ_head_. When analyzing the mean improvement in decoding performance achieved beyond all other predictors, we found that this value was larger for position (0.048) than for θ_head_ (0.019; all p < 10^−15^, Wilcoxon’s signed-rank tests) or the other two covariates. Furthermore, the improvement due to θ_head_ was larger than for these other factors (speed: 0.011; p < 10^−15^; Δθ_head_: 0.008; p < 10^−15^, Wilcoxon’s signed-rank tests).

Both the conditional mutual information and encoding analysis show that neurons can carry significant information on both factors. However, other factors such as stimulus availability, choice and reward may have influenced these results, because during track running they are closely associated with both position and head direction in space and/or time (Zaidel et al., 2017). Re-analysis of conditional information, now on data in which the period of stimulus presentation ± 1 s was eliminated, showed a decrease in the amount of information on θ_head_ conditional on position, which was nevertheless still significantly larger than 0 for both areas (cMI difference from 0, A1: p = 1.7 ⁎ 10^−4^, V2L: p < 10^−15^, Wilcoxon signed rank tests; Supplementary Figure 2A, cf. Fig. 4B). Similarly, re-analysis of encoding improvement with data from which the stimulus presentation period ± 1 s was eliminated yielded similar results compared with the original finding (Supplementary Fig. 2B, cf. Fig. 4G). Together, these results suggest that position and, to a lesser degree, head direction are important drivers of firing activity of neurons in A1 and V2L. However, we cannot rule out potential contributions to information related to position or head direction as a result of correlations with additional subject-induced sensations (e.g. eye movements), which we were unable to detect in the current dataset.

### Coherent mapping of location in A1 and V2L populations

If the neuronal populations in A1 and V2L code the animal’s track position, it should be possible to infer the position of the rat from the collective neuronal activity. We used a Bayesian decoder to test whether spatial location can indeed be predicted from population activity (sessions with >16 neurons/area were included). Decoding of position was successful for both A1 and V2L, because the chance that a decoded position matched the true position of the rat was larger than the chance that it matched any other location on the track (i.e. clear diagonal structure in Fig. 5A, B; A1: n = 15 sessions; V2L: n = 12 sessions). The distributions of decoding errors, which are the Euclidean distances between the true and decoded positions, clustered near 0 m for both areas (Fig. 5C). V2L showed, on average, higher proportions of smaller errors than A1, indicating that decoding spatial position from V2L was more accurate than from A1 (V2L error: 0.11, 0.10 - 0.42; A1 error: 0.29, 0.26 - 0.37; median and interquartile range, in meters).

**Figure 5:**
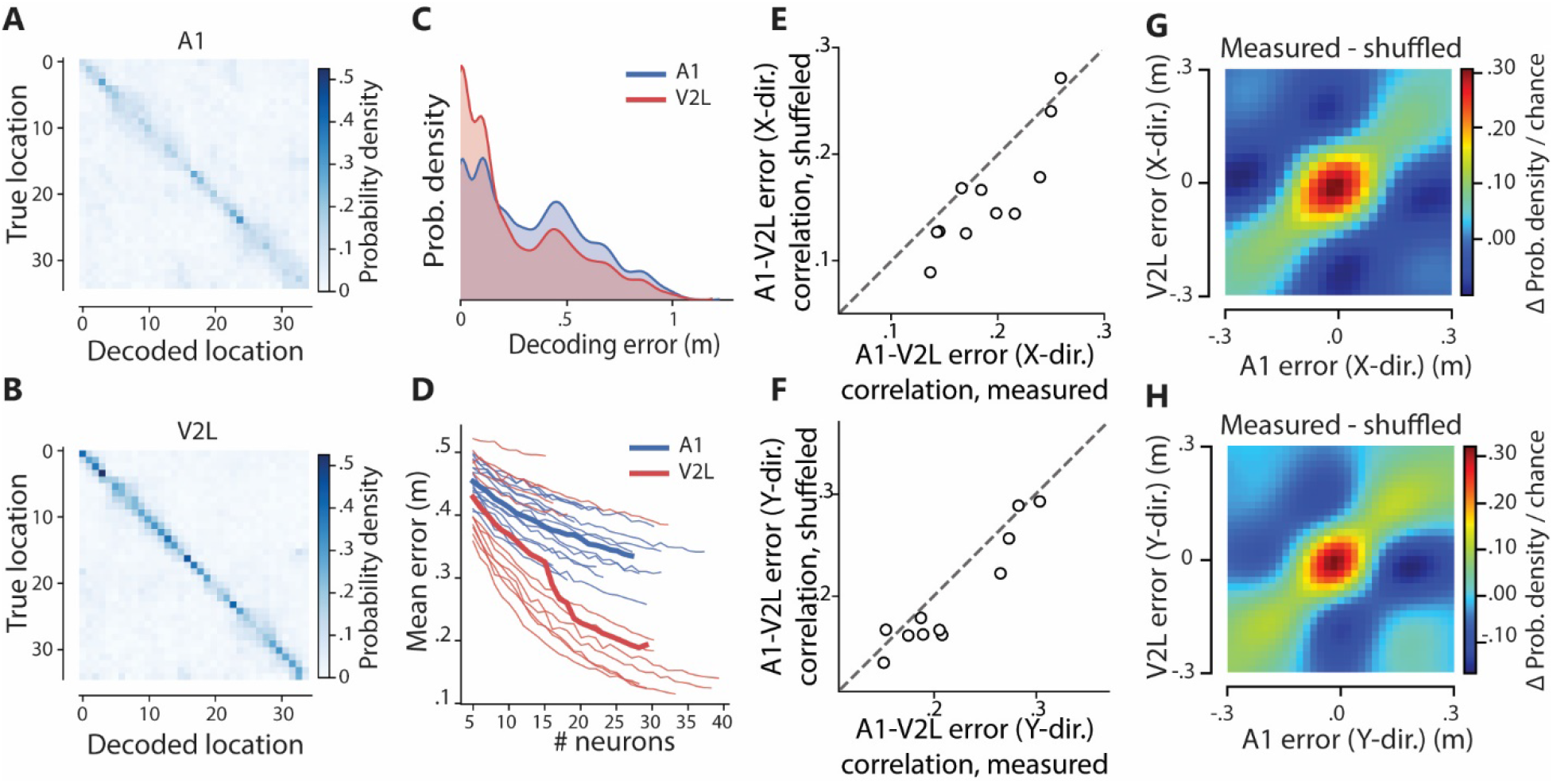
Spatial position can be reconstructed from A1 and V2L populations using a Bayesian decoder. **(A)** Confusion matrices indicating the performance of Bayesian decoding of linearized position from A1 neuronal populations averaged across sessions. The probability that a sample of spiking activity is assigned by the decoder to the true location of the rat is coded in blue shades. **(B)** as (A) but for V2L populations. **(C)** Distribution of decoding errors (i.e. distances between the true and decoded position) across A1 (blue) and V2L (red) sessions. **(D)** Decoding error as a function of population size, obtained by randomly selecting units from the neuronal populations. Thin lines indicate means for individual sessions for A1 (blue) and V2L (red). Thick lines indicate means over sessions. **(E)** Correlations of measured (abscissa) and shuffled (ordinate) instantaneous decoding errors in the X-dimension of the maze. Shuffling was performed inside the same spatial bin and within the same running speed range. **(F)** Same as (E) but for the Y-dimension. **(G)** The size and direction of instantaneous errors were correlated between A1 and V2L. Color coded is the frequency of observing an instantaneous decoding error of a particular size and direction in the X-dimension after subtracting the frequency of instantaneous errors of a particular size and direction following shuffling of the errors within the same spatial bin and speed range. E.g. a value of +0.3 indicates that an error of that particular size and direction is 30% more likely to be observed in actual data than in shuffled data. **(H)** Same as (G) but for the Y-dimension.

Increasing the number of neurons included in the decoding analysis decreased the mean decoding error for both areas. We considered several population sizes, which were similar for both areas (A1: 16- 38 neurons, V2L: 16-40 neurons), but bootstrap analysis showed no signs of plateau performance with progressive population sizes (Fig. 5D). Although decoding from both areas likely could be improved had more neurons been recorded, at identical population size decoding performance for V2L was better than for A1, both for the mean across sessions and for most individual sessions.

To investigate whether locations are encoded coherently across both areas, we calculated the correlation in instantaneous decoding error between A1 and V2L. Instead of only taking the magnitude of the error into account, errors in the *x* and *y* dimensions of the maze were considered separately and their directionality was preserved. When decoding errors were computed for sessions which contained >16 simultaneously recorded neurons in each area, we found significant correlations (p < 0.001 for all 11 sessions). This was also the case when including all sessions where at least one of the two areas provided 16 neurons, again indicating that A1 and V2L encode locations coherently. However, these correlations may be confounded if certain locations are represented more accurately than others in both areas, or if firing patterns are subjected to a common modulation by locomotion speed. To control for these possibilities, we shuffled the data across time points at which the animal was in the same spatial bin and running within the same range of speeds. The correlations between the shuffled decoding errors were significantly lower than the observed correlations (x-direction, p = 0.01, y-direction p = 0.02, Wilcoxon signed-rank test). This difference remained significant when we included all sessions where at least one area provided 16 neurons (x-direction, p = 0.001, y-direction, p = 0.01). The residual decoding errors, obtained by subtracting the shuffled distribution from the observed joint distributions, displayed a diagonal structure (Fig. 5G-H), indicating that representations of location in A1 and V2L remain coherent even when position cannot be decoded accurately from the population of either area. This coherency exceeds what would be expected from a common influence of speed. In addition to the mean across sessions (Fig. 5G-H), these results also held for individual rats (Supplementary Figure S3). It is unlikely that error correlations can be accounted for by decoding artifacts in sessions with poor decoding performance because there was no evidence of a positive relationship between instantaneous decoding error correlations and average decoding error, rather their linear correlations were negative and not significant (Supplementary Fig. 4).

## Discussion

We showed location- and head direction-selective firing patterns in large proportions of neurons in A1 and V2L of freely moving rats navigating a figure-8 maze. This activity takes part in coherent representations across areas, as indicated by highly correlated firing-field densities (fig. 3A-C), correlated errors in reconstructed position (fig. 5) and cross-areal noise correlations which varied as a function of spatial tuning and immediate position (fig. 3D-F).

A first implication is that neural representations bound to subject location exist in the sensory cortex outside the visual system. Such activity may be expected in V2L because of similar observations in V1 (Fiser et al., 2016; Fournier et al., 2019; Haggerty and Ji, 2015; Ji and Wilson, 2007; Saleem et al., 2018) and egocentric trajectory correlates in spiking activity of posterior parietal cortex (Krumin et al., 2018; McNaughton et al., 1994; Nitz, 2006; Whitlock et al., 2012). These areas share anatomical borders with V2L and bidirectional, monosynaptic connectivity (Haggerty and Ji, 2015; Krumin et al., 2018; McNaughton et al., 1994; Miller and Vogt, 1984). That A1 neurons also show location-tuning is more surprising, because A1 activity has hitherto not been associated with animal location. However, in many mammals this area is required for sound localization (Heffner, 1978; Jenkins and Merzenich, 1984; Kavanagh and Kelly, 1987; Thompson and Cortez, 1983) and contains neurons tuned to the location of sound sources (Middlebrooks and Pettigrew, 1981; Town et al., 2017; Wang et al., 2019).

### Nature and function of location-selective firing in sensory cortices

In addition to single or multiple peaks in single-cell firing rates tessellating the entire maze, we found that A1 and V2L spike patterns were spatially stable across trials and coded significant amounts of information on animal position (fig. 2 and 3). Our maze harbored repetitive elements requiring the same local behavior, such as directional body turns or running along straight maze stretches, whereas many cells showed single firing-rate peaks and thus did not reflect these repetitions (fig. 2). Furthermore, the encoding analysis revealed animal position as the strongest predictor of A1 and V2L activity, even after correcting for head direction (fig. 4). Moreover, subject position could be inferred from A1 and V2L ensemble activity using Bayesian decoding (fig. 5). Despite these indications, we argue that neither our current findings, nor previous results on V1 (Fiser et al., 2016; Haggerty and Ji, 2015; Ji and Wilson, 2007; Saleem et al., 2018), necessarily imply coding of (allocentric) position *per se*, because proving this would require additional experimental manipulations to establish that location-selective activity is independent of the locomotion direction through a location and tolerates manipulation of local sensory cues (Knierim, 2002; Lansink et al., 2012; Leutgeb et al., 2005; Speakman and O’Keefe, 1990; Wilber et al., 2014).

We propose the more general alternative that sensory cortical areas integrate modality-specific evidence with information from other sensory, motor and association areas to generate sequential representations, which are not merely sensitive to local sensorimotor cues, but also to contextual elements (which may comprise spatial but also other task-relevant elements such as reward proximity, task rule execution, etc.). For instance, activation of visual cortex neurons depends on the subject’s field of view, which in turn depends on animal position and head-direction (cf. Haggerty and Ji, 2015). This combination of view, position and head direction may give rise to a predictive visual representation in the cortex, which will be compared to further visual input to compute error signals, as posited by predictive processing models (Fiser et al., 2016; Friston, 2005; Keller and Mrsic-Flogel, 2018; Pennartz et al., 2019; Rao and Ballard, 1999). Although this proposal needs further testing, it is generally supported by the literature documenting extensive corticocortical connections (Bizley et al., 2007; Budinger and Scheich, 2009; D’Souza et al., 2016; Felleman and Van Essen, 1991; Gămănuţ et al., 2018; Harris et al., 2019; Laramée et al., 2011; Leinweber et al., 2017), contributions to neural coding in sensory cortices by non-sensory parameters (Goltstein et al., 2013; Namboodiri et al., 2015; Pakan et al., 2018; Shuler and Bear, 2006) and auditory-visual cortical interactions (Ibrahim et al., 2016; Iurilli et al., 2012; Knöpfel et al., 2019; Meijer et al., 2020; Meijer et al., 2017; Morrill and Hasenstaub, 2018).

This view does not conflict with a potential role for sensory cortices in updating spatial (e.g. hippocampal) representations in a bottom-up fashion, or with the navigational system contributing to top-down sensory predictions (Fournier et al., 2020). However, the hypothesis of the hippocampus causally driving spatial coding in V1 (Saleem et al., 2018) faces the issue that the hippocampus proper does not directly project to the sensory cortices, and its output is transformed by the synaptic matrices of intermediate parahippocampal regions on which also non-hippocampal structures converge (Furtak et al., 2007; Rusu and Pennartz, 2020; Witter et al., 2000). Instead of guiding spatial navigation, the bidirectional, cortico-hippocampal circuitry may subserve declarative memory consolidation (Eichenbaum, 2000; McGaugh, 2000; O’Keefe and Nadel, 1978; Rusu and Pennartz, 2020; Squire, 1986). In this process, distributed neocortical activity selective for parts of a behavioral sequence may form a common ‘pointer’ or marker for binding together cross-modal information (Teyler and DiScenna, 1985).

### Sensitivity to head direction in A1 and V2L

For 33% of neurons in A1 and 49% in V2L, the information conveyed on head direction was significant after correcting for position (fig. 4B, 4D). As for subject location, this finding relates to how sensory-specific states coded by A1 and V2L depend on head direction, or how these sensory cortices may use this sensitivity to emit head direction signals to target areas. Head direction signaling is regulated by the vestibular nuclei carrying information about the head’s motion relative to external space (Cullen, 2014) and, in rodents, about static neck position (Barresi et al., 2013; Medrea and Cullen, 2013). Vestibular input is a key contributor to the brain’s head-direction system (Taube, 2007), including anterior thalamus (Taube, 1995), post-subiculum (Taube et al., 1990) and medial entorhinal cortex (Sargolini et al., 2006). Vestibular information was shown to reach the visual system directly and indirectly, viz. via the retrosplenial cortex (Vélez-Fort et al., 2018), which was proposed to mediate between the sensory cortices and the head direction system of the temporal lobe (Page and Jeffery, 2018). Whether head-direction signaling in A1 and V2L has a causal role in distributing head-direction information to target areas is up for further research. The collective evidence supports the hypothesis that areas along the cortical hierarchy may use both allocentric and egocentric representations, with a gradient of egocentric-to-allocentric processing from sensory to temporal cortices, as also proposed for parietal-retrosplenial circuitry, where V2L is sometimes included as part of parietal cortex (Chen et al., 2018; Clark et al., 2018; Wilber et al., 2014).

### Comparison between primary auditory cortex and secondary visual cortex

We were particularly struck by the broad, qualitative similarities between A1 and V2L spiking patterns. Both areas showed comparable levels of spatial stability, amounts of firing fields per unit (fig. 2), spatial distributions of firing fields and noise correlations (fig. 3). Moreover, the contributions to predictions of firing-rate patterns from position, head direction and other factors were highly similar for A1 and V2L (fig. 4). These observations lend support to the hypothesis that coordinated representations are a general feature of sensory cortical areas.

Neural coding in A1 and V2L also showed interesting quantitative differences, which consistently point to a higher spatial information content (fig. 2), stronger correlations with location and head direction (fig. 4) and better position reconstruction in V2L than A1. How this greater accuracy arises in V2L is unknown, but it may relate to a larger amount of spatially informative visual cues in our maze compared to auditory cues, and to more consistent changes in visual inputs due to self motion than to (self-induced) auditory inputs. In view of this greater accuracy, it is interesting to observe the slightly earlier firing in A1 than V2L units (fig. 3G). This might reflect the exploitation of potentially faster, subcortical processing of auditory compared to visual information (Picton et al., 1974).

### Significance of cross-areal coordination in cortical mapping of location and head direction

Arguably our most novel result is that location-selective representations are highly coherent across sensory domains of the cortex (fig. 3, 5), suggesting that location- and head direction-sensitive mappings in auditory and visual cortical systems are not computed independently but are coordinated. Such coordination of context-dependent sensory mappings offers the computational advantage that evidence from multiple sensory modalities can be combined to improve estimates of the subject’s task-relevant state, although this comes at the expense of error sharing.

Previous studies on auditory-visual interactions were predominantly guided by the theoretical framework of multisensory cue integration, whereby evidence for the detection of a stimulus in one modality is augmented by another modality (Fetsch et al., 2013; Meijer et al., 2019; Stein et al., 2014). Our findings go beyond integration of discrete sensory cues and instantaneous sensory states as they indicate a cross-modal coordination of context-sensitive representations. In our proposal this common mapping subserves the construction of a multimodal survey of the subject’s current situation, thereby enabling efficient goal-directed action planning and execution (Pennartz, 2018). As argued by Hawkins et al. (2017), self-parameters including head direction and position within a task sequence are key priors in determining how and when to undertake goal-directed actions. In line with predictive processing, knowledge of these self-parameters is required to interpret novel sensory inputs and anticipate sensory outcomes of actions (cf. Schürmann et al., 2019).

## Materials and Methods

### Experimental design

#### Subjects

Experiments were performed on Lister Hooded rats (n=3, Envigo, the Netherlands) at an age between 9 and 40 weeks. All rats were socially housed during behavioral training, but individually housed during periods of recordings when rats had an implanted tetrode-array, under a normal day/night cycle (lights on: 8:00am, lights off: 8:00pm). The rat’s food intake was restricted such that its weight was at least 85% relative to the standard growth curve provided by the breeder (Envigo, the Netherlands), corrected for the deviance in weight between the rat and the curve in the week before the start of food restriction. Weights were maintained at a stable level after rats reached healthy, adult body weight (Clemens et al., 2014; Newby et al., 1990). Rats had *ad libitum* access to water throughout the experiment. All experiments were performed in accordance with the National Guidelines on Animal Experiments and were approved by the Animal Experimentation Committee of the University of Amsterdam.

#### Behavioral setup

Rats were trained to discriminate between auditory and/or visual stimuli on an automated, rectangular figure-8 shaped track (92 cm x 73 cm) which was raised 55 cm off the ground (fig. 1A). The track’s alleys were made of black painted aluminum (width = 8.7 cm) and contained raised edges (1.0 cm). At the front of the track, two LCD monitors (Iiyama ProLite B2776HDS) and two audio speakers (Audaphon Neo CD 3.0) were available for stimulus presentation. The monitors were positioned symmetrically from the center of the track and the speakers were located above the top-left and top-right corners of the left and right monitors, respectively. During early training stages, transparent polycarbonate walls lined the central alley to prevent the rat from prematurely exiting this alley. Additionally, two transparent polycarbonate sliding doors were positioned at the front and back of the central alley. The walls and door at the front of the central alley contained small holes to allow for perception of the auditory stimuli. Two reward wells were positioned at the left and right edges of the track’s front alley. An additional reward well was positioned in the central alley, towards the front-end T-junction. Fluid sucrose solution (15% in tap water) was delivered to the wells by syringe pumps (Razel, VT, USA). All reward wells contained infrared photodetectors to detect nose pokes and licks. The motor activity of the rat on the track was registered with additional photodetectors, located near the T-junctions at the front and back ends.

The track was entirely computer controlled, obviating the need for human interventions during the experiment, and was interfaced with the recording system to ensure synchronized time stamping of behavioral events and neuronal activity patterns. It was positioned in an enclosure of black curtains (2.8 x 2.2 m) within a sound-attenuated room of 3 x 3 m. The room was dimly lit by a small LED light pointed towards the ceiling. The experimenter observed training and experiments from an adjacent room to minimize interference with the rats’ behavior.

#### Stimuli

On each trial of the behavioral task, rats had to discriminate between a target stimulus, displayed on one of the screens and/or the adjacent speaker, and a distractor stimulus, appearing on the other screen and/or speaker (Fig. 1B). Target and distractor were either unisensory (visual or auditory) or multisensory (audiovisual) and differed in stimulus amplitude. Three types of target/distractor combinations were in equal proportions presented to the rat; large-difference unisensory trials (1/3), small difference unisensory trials (1/3) and small difference cross-modal trials (1/3). In a separate session for each rat, the threshold amplitude differences (i.e. the amplitude differences at which the rat shows a correct response in 50% of the stimulus presentations) were determined using a staircase procedure. On the basis of this data, stimulus parameters were set for each rat such that the discrimination performance for the large-difference unisensory trials was >70% correct and for the small-difference unisensory trials was >50% correct. The stimulus settings remained the same for all recording sessions of a rat. The amplitude differences for the stimulus components in the audiovisual trials were identical to the differences for the small-difference unimodal stimuli of the same session. The specific screens (left or right) at which target and distractor were displayed were pseudorandomly selected for each trial, such that the target stimulus was never displayed more than four successive trials at the same side and the difference in left and right target-presentations across the session was not more than 4.

Visual stimuli were full-screen, upward-moving, black-and-white checkerboards (0.1 cycles/degree, 4 cycles/second; fig. 1B). When no visual stimuli were displayed, a grey background was visible. All visual stimuli and the background were gamma corrected and had the same overall luminance as measured with a photometer. Contrast values between light and dark checkers varied between 0 and 1, in which 0 indicates no contrast and 1 indicates the maximum contrast possible with the monitor at its lowest brightness setting.

Auditory stimuli were composed of white noise, which was band-passed between 10 and 25 KHz. When auditory stimuli were absent, background noise was played. Background noise was white noise band-passed between 8 and 12 KHz. For auditory stimuli, the difference between target and distractor was in the relative volume between the speakers. Relative auditory volume ranges between values of 0 and 1, with 0 and 1 indicating that the volume is fully accounted for by one or the other speaker. Background noise was played with a contrast of 0.5, i.e. identical volume through both speakers. Therefore, at every moment during the session, sound was playing at the same volume, which was set to 76 dB at the central reward well.

#### Behavioral training

After rats had learned to complete unidirectional laps on the track and to nose-poke for a duration of 1 second to earn reward, they were trained to discriminate between target and distractor stimuli and to respond by choosing the side at which the target stimulus was present. When the front door opened at the start of the trial, the target visual stimulus was presented on one screen (i.e. on one side of the track) while the other screen maintained the grey background. The rat earned a reward if it poked in the well at the side of the track corresponding to where the stimulus was displayed. The stimulus presentation lasted until 3 seconds following reward delivery, with the aim of strengthening the stimulus-response-outcome association. When the rat made a nose poke at the incorrect side, stimulus presentation stopped immediately. From this stage onwards, the length of the nose pokes to start stimulus presentation was increased incrementally from 0 to 0.5-1.5 seconds (randomized across trials). If the rat performed at least 60 trials within 60 minutes with 70% correct trials on three out of five consecutive days, stimuli switched from visual to auditory. If this criterion was met also for auditory stimuli, subsequent sessions included audio, visual stimuli and audio-visual stimuli in equal proportions and presented according to the pseudorandom schedule (see *Stimuli*). In multisensory trials, the auditory and visual stimulus components were always presented on the same side of the track; i.e. no sensory conflicts were created. Once the rat reached the same criterion with these three trials types included in the session, the discrimination problem was made progressively more difficult, by introducing distractor stimuli and lowering the contrast of visual target stimuli in subsequent sessions. The difference in amplitude between target and distractor stimuli was gradually decreased over training sessions but never below the level described above for ‘low-difference stimuli’. In parallel with the increase in difficulty, the display time of the stimuli was progressively shortened until the stimulus duration was 2-3 s. To prevent rats from developing habitual or stereotyped response preferences, extra sessions were occasionally included in the training. In these sessions, rats were allowed to collect reward from the correct well after sampling the incorrect well. After an incorrect nose poke, trials continued until the correct well was sampled. These sessions did not count towards criterion performance.

In the final task, the trial procedure was similar to training stage 4 and now the small and large difference unimodal stimuli were presented alongside with the crossmodal stimuli (see *Stimuli)*. To ensure enough trials were performed in all conditions to allow statistical comparison, each session contained either visual or auditory unimodal trials, in addition to multisensory trials. Recordings commenced when rats consistently performed at criterion level: >60 trials in 60 minutes with >70% correct on large-difference unimodal stimuli and above chance level for small difference unimodal stimuli and multisensory stimuli.

#### Tetrode array and surgical procedure

A custom-made, 128 channel tetrode-array was implanted over the right hemisphere of three rats (Bos et al., 2017; Lansink et al., 2007). Four bundles of 8 individually moveable tetrodes each were targeted to primary auditory cortex (A1, −4.4 mm AP, 6.8 mm ML, −4.4 mm DV), lateral secondary visual cortex (V2L, − 5.8 mm AP, 6.0 mm ML, −2.6 mm DV), hippocampal area CA1 (CA1, −3.6 mm AP, 2.4 mm ML, −2.6 mm DV) and perirhinal cortex area 36 (PRH, −4.4 mm AP, 6.8 mm ML, −7.0 mm DV). Hippocampal and perirhinal data were not used in the current study. Thirty minutes before surgery, rats received the analgesics meloxicam (Metacam, 2mg/kg) and buprenorphine (Buprecare, 0.04 mg/kg) subcutaneously, as well as the antibiotic enrofloxacin (Baytril, 5mg/kg). Anesthesia was induced using 3% isoflurane in oxygen and maintained with 1-2% isoflurane. Animals were mounted in a stereotaxic device (Kopf; Tujunga, CA, USA) and placed on a heating pad to maintain their body temperature. A single craniotomy was made such that the points at which the bundles entered the brain were positioned relative to bregma at −5.4 mm AP and 2.8 mm ML (A1), −5.9 mm AP and 4.0 mm ML (V2L), −3.6 mm AP and 1.7 mm ML (CA1) and −3.5 mm AP and 3.8 mm ML (PRH). Six screws were placed into the skull, with the screw positioned over the frontal bone serving as electrical ground for the tetrode array. The hyperdrive was positioned such that the bottom of the bundles touched the cortical surface. The craniotomy was then sealed using silicone adhesive (Kwik-Sil) and dental cement was used to fix the hyperdrive and screws to the skull. Post-operative care consisted of subcutaneous injections of the analgesic meloxicam (2 days) and Baytril antibiotic (1 day) and application of wound healing ointment (Acederm, Ecuphar, Breda, Netherlands). On post-operative days 1-3, rats received 10 g of extra food softened in water to facilitate consumption. Tetrodes were gradually lowered to their target regions across the first week after surgery. During recordings their depth was estimated from the number of turns to the guiding screws and from the online Local Field Potential (LFP) profiles.

#### Data acquisition and preprocessing

Spikes and LFPs were recorded using tetrodes (Gray et al., 1995)(nichrome, California Fine Wire, 16 µ per lead, gold-plated to an impedance of 500-800 KOhm) using a Neuralynx Digitalynx SX recording system (Neuralynx, Bozeman MT). Raw signals were buffered using four 32-channel unity-gain head stage amplifiers before being passed through an automated commutator (Neuralynx, MN). Each of the four tetrode bundles contained an additional electrode that served as a reference channel and which was positioned in the white matter near the tetrode bundle. The recorded signals were the raw signals with the reference signal subtracted. For spike recordings, signals were band pass filtered between 600 and 6000 Hz. Putative spikes were recorded for 1 ms (16 bit, 32KHz) from all leads of a tetrode whenever the signal on any lead of that tetrode crossed a predefined threshold. LFPs were low-pass filtered below 300 Hz and recorded continuously (16 bit, 3.2 KHz). The behavior of the rat was tracked by a ceiling-mounted camera (at 720 by 576 pixels, 25 fps) and timestamped by the Digitalynx SX.

Spikes were attributed off-line to putative single units (clusters) using the KlustaKwik automatic clustering algorithm (Kadir et al., 2014) followed by manual refinement (MClust 3.5). Waveform features used for clustering were energy, the first derivative of the energy, the overall peak height and the peak height during samples 6-11 where the action potential peak is expected. Clusters were included for further analysis based on a combination of quality metrics (Schmitzer-Torbert et al., 2005): L-ratio (< 0.2 – 0.8), isolation distance (> 15 – 24) and interspike interval (1.2ms) violations (< 0.1% – 0.5%).

#### Histology

Following the last recording session animals were anesthetized using Isoflurane and electrolytic lesions were made at each tetrode tip by passing current (18 uA for 2s) through two leads of each tetrode. At least twenty-four hours later, the animal was deeply anesthetized with an intraperitoneal injection of Nembutal (1.0 ml, sodium pentobarbital, 60 mg/ml, Ceva Sante Animale, Maassluis, Netherlands) and transcardially perfused with 0.9% NaCl solution, followed by perfusion of a 4% paraformaldehyde solution (pH 7.4 phosphate buffered). The brain was extracted and placed in 4% paraformaldehyde solution for at least 24 hours post-fixation, after which 40 um transverse sections were made using a vibratome. These were stained with Cresyl Violet which allowed reconstruction of tetrode tracks and their endpoints marked by electrolytic lesions (Paxinos and Watson, 2007)(Supplementary fig. 1). Because it was not possible to determine with absolute certainty how each endpoint corresponded to the identity of individual tetrodes, the considered neuronal pools likely contain a small subset of units recorded in the vicinity of the target locations. Based on the average number of neurons included from the respective recording sessions for A1 and V2L, the estimated number of neurons recorded outside their target areas was 16 for V2L and 32 for A1.

### Statistical Analysis

#### Statistical procedures

Unless specified otherwise all statistics were performed using linear mixed models (LMMs) or generalized linear mixed models (GLMMs) in MatLab (MathWorks, Natick, MA), depending on the distribution of the data. All reported statistical quantities (group means, regression slopes, confidence intervals etc.) were derived from the (G)LMMs. All reported confidence intervals are 95% prediction intervals. To estimate the denominator degrees of freedom (DF2) for F-tests, the Satterthwaite approximation was used for LMMs and the residual degrees of freedom for GLMMs. To correct for multiple comparisons, p-values are adjusted using the Holm-Bonferroni method where applicable {Holm, 1979 #172}(fig. 2H, 3E-F).

#### Inclusion criteria

Behavioral performance was measured as the percentage of correct responses to stimulus presentation in a session. Sessions were included for analysis only if the performance was above 95% of a distribution of performance expected from sessions with an equal number of trials and chance performance (50% correct trials). Additionally, to exclude sessions where animals had a preference for a particular response direction, sessions were included only when responses to both left and right stimuli were performed above chance using the same procedure.

#### Z-scored spiking activity and firing rates

Spike trains of individual units were binned into 1 ms temporal bins and smoothed with an exponential window with a time constant of 150 ms. Z-scored spiking activity was calculated by subtracting the mean from the smoothed spike train and dividing the result by the standard deviation.

#### Position tracking and 2D rate maps

The location of the rat’s body and head were determined separately for each frame of recorded video using custom scripts made with Bonsai Editor (Lopes et al., 2015). Frames with erroneously assigned positions were manually corrected. The raw tracking data was smoothed using the *smooth()* function in Matlab using the ‘*rlowess*’ method and a span of 5 pixels. Head direction was determined for each video frame as the angle of the body-head axis of the rat with respect to the left-to-right axis of the setup.

Rate maps were constructed in 2D by spatially binning (bin size = 10 by 10 pixels or 4.2cm^2^) the smoothed spike trains of single units into the spatial bin occupied by the rat in each videoframe, summing the firing rates per bin and then averaging over the time spent in each bin (occupancy). Binned spike trains and occupancy maps were independently smoothed by convolution with a 2D Gaussian (std = 1 bin) before averaging. Time segments in which the running velocity of the rat was < 6 cm/s were excluded.

#### Linearization of position data and 1D rate maps

The goal of linearizing position data is to allow more powerful analyses relating localized spiking activity to behavior. The lateral range of body motion on the track was limited, and the firing fields observed on the 2D rate maps generally spanned the full width of the track. Linearization therefore allows to focus on the spatial dimension containing the majority of the rate map structure. Linearization of the rat’s position was achieved by first determining the average path of the rat across the setup for each recording session. This was achieved by manually tracing the locations with highest occupancy of each session’s occupancy map. The starting point of the linearized track was chosen as the starting point of the right alley segment at the three-way junction at the front of the setup (FC; fig. 1A). The linearized track then consisted, in this order, of the right side of the track, central alley, left side of the track. The end position was the center of the back-alley segment at the three-way junction. The linear position was then determined as the point along this linearized track closest to each observed 2D position. Raw linearized position was smoothed by convolution with a Gaussian with standard deviation of 2 pixels (0.4 cm). Linear speed was then calculated from the linear location using the *gradient()* function in MatLab, and linear acceleration was calculated similarly from linear speed. Finally, because the linearized track length varied slightly between rats, linear location was normalized to the mean length across animals (334 cm). Linear rate maps were made similarly to the 2D rate maps but using the linear position data, using the same inclusion criteria and smoothing parameters and with a bin size of 16 pixels (3.3 cm). Joint rate maps (fig. 3A-B) were constructed from the linear rate maps of each neuron by normalizing each rate map between 0 and 1, and then sorting all rate maps of all sessions and rats by the location of the peak firing rate (dark red in fig. 3A).

#### Firing fields

The procedure for determining the location of firing fields was similar to the method described by (Haggerty and Ji, 2015), using the unit’s linear rate map as a basis. First the rate map was smoothed using the *smooth()* function in Matlab (MathWorks, Natick, MA) with the ‘*rlowess*’ method and a span of 5 bins. The baseline firing rate was determined as the 40^th^ percentile of this smoothed rate map. The baseline was subtracted from the rate map and local maxima were determined for the baseline-corrected rate map. Local maxima were kept for further processing if the rate was > 1Hz and > 0.2x the baseline rate. Field boundaries were determined as the bins nearest to a peak where the firing rate was < 10% of the peak firing rate. This procedure sometimes produced very small fields near or on the slope of larger fields (‘shoulders’). Such shoulders were discarded if the border of the field was two or fewer bins apart (<=6.6cm) from the border of a taller field, unless it was two or more bins (>=6.6cm) away from its own border and the peak was > 1.5 times the value at this border. After the removal of spurious small fields, the borders of remaining fields were extended to the bins where the activity fell below the peak cut-off of 1 Hz and 0.2 times the baseline rate, or until they reached the border of another peak. This procedure was followed for each single unit from the tallest to the lowest peak. Single fields that spanned the T-junctions at the front and back of the experimental track were prevented from being detected as two separate fields by considering the bins of the linear rate maps that border the T-junctions as adjacent.

#### Spatial Stability

The spatial stability of the localized firing of individual units was determined using a permutation analysis. First, separate linear rate maps of each individual trial were constructed, similarly to the procedure described above, except that the data was split into leftward and rightward trials. The correlation between a single-trial rate map and the rate maps of all other trials on the same side of the track was computed (Pearson correlation), for each trial and then the average correlation coefficient was computed. These steps were repeated for 1000 shuffled versions of the spiking data. For each shuffled iteration, the spiking data was temporally rotated within a single trial by a random number of samples (Louie and Wilson, 2001). This method of shuffling leaves the temporal structure of spiking patterns largely intact. Positions and spikes emitted during periods of immobility (<0.06 cm/s) were excluded from analysis. A unit was considered spatially stable if its average single-trial rate map correlation, for both left- and rightward laps, was higher than 95% of the distribution of average, shuffled single-trial rate map correlations of the corresponding side. To calculate the spatial stability index, the mean of the distribution of average, shuffled single-trial rate map correlations was subtracted from the average observed single-trial rate map correlation; the result was divided by the standard deviation of the shuffled distribution. This procedure was done for each unit for both left- and rightward trials and the reported spatial stability index was the mean of those two.

The probability of the spatial stability of a unit was modeled using a GLMM with link function *g*(*P_stable_*) = ln (*P_stable_*) and the equation:

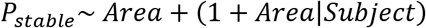

Where *Area* is a categorical variable indicating the cortical area where the unit was recorded and *Subject* is a categorical variable indicating from which experimental subject the data originated.

#### Spatial information

Spatial information (SI) for each unit was calculated from the linearized location data and firing rate following Skaggs et al. (1992):

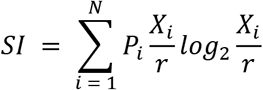

Where *P_i_* is the probability of finding the animal in bin *i*, *X_i_* is the sum of the firing rates observed when the animal was found in bin *i*, *r* is the mean spiking activity of the neuron and *N* is the number of bins of the linearized trajectory (104). The SI for each unit was modeled with a GLMM using a gamma distribution with link function *g*(*SI*) = *SI*^−0.01^ and the equation:

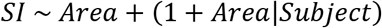

#### Correlating firing field densities

Firing field densities were calculated per rat by first determining for each linear spatial bin the number of firing fields, across all units, in which this bin took part. Then this number was divided by the subject’s total number of fields. The average firing field density is reported as the mean of the field densities for the individual rats. Firing field densities of A1 and V2L were correlated with an LMM using the equation:

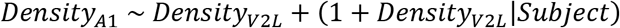

#### Linear regression of spiking activity on behavioral covariates

Linear regression was used to determine whether firing field correlations between A1 and V2L could be explained by similar linear dependencies between single unit firing rates and behavioral covariates. The instantaneous, z-scored spiking activity of each unit was regressed on linear running speed, acceleration, head direction and change in head direction (angular velocity) using the general linear model:

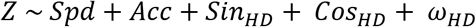

Where *Z* is the instantaneous, z-scored spiking activity, *Spd* is linear speed, *Acc* is linear acceleration, *Sin_HD_* and *Cos_HD_* are the sine and cosine of head direction and *ω_HD_* is the angular velocity in head direction. The model residuals of each unit were used to produce linear rate maps and subsequently determine firing fields without linear dependencies on the behavioral covariates.

#### Stimulus responsiveness

The responsiveness of a unit to the sensory stimuli was assessed by statistically comparing the mean, z-scored firing rates of the time intervals of [350 – 50] ms pre-stimulus onset and [0 – 300] ms post-stimulus onset using Wilcoxon’s signed-rank test (alpha = 0.05). A subset of units gradually increased or decreased firing rates before stimulus onset (“ramping activity”), without showing a change in spiking activity at stimulus onset. To preclude that such activity would erroneously be considered as significantly stimulus responsive, only units with a stable pre-stimulus onset firing rate were considered; i.e. the unit’s firing rates between [1000-700 ms] and [350-50] ms pre-stimulus onset were required to be similar (Wilcoxon signed rank test, P > 0.05). The probability of a unit’s responsiveness to stimuli was modelled using a *g*(*P_resp_*) = ln(*P_resp_*)and equation:

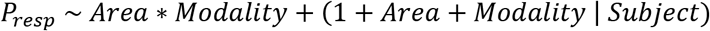

Where *Modality* is a categorical variable indicating the modality (auditory or visual) of the stimulus being considered.

#### Noise correlations

Rate map similarity between neurons was first determined for each pair of units as the Pearson correlation coefficient of their linear rate maps. To determine noise correlations, we first temporally binned the instantaneous, z-scored spiking activity into 10 ms bins and then subtracted from that the mean spiking activity at the linear spatial bin occupied by the rat at each instant. This difference constitutes instantaneous variability in spiking activity not predicted by spatial position. Additionally, for each pair of units we categorized locations along the track as i) occurring in a firing field of both units (“shared field bin”), ii) occurring in a firing field of only one of the units (“one-field bin”) or iii) occurring outside of firing fields of both units (“out of field bin”). For each pair of units we calculated noise correlations as the Pearson correlation coefficient of the instantaneous variability of the activity of two units, and we did so separately for each of the three bin types.

Analysis proceeded with pairs showing significant noise correlations, determined via comparison to shuffled distributions of these correlations. These distributions were made by computing noise correlations after shuffling samples of spiking activity within the same linear spatial bin and repeating this procedure 5000 times. A pair was considered significantly correlated if its actual correlation exceeded 99% of the shuffled distribution for at least one of the three location types.

For significantly correlated pairs, mean noise correlations across location types, were regressed on their rate map correlations with an LMM using the equation:

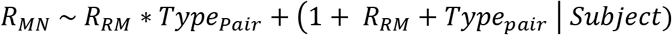

Where *R_MN_* is the mean noise correlation of a pair across location types, *R_RM_* is the rate map correlation and Type*_pair_* is the type of pair (A1-A1, V2L-V2L or A1-V2L). Here we used the mean noise correlation across location types rather than the overall noise correlation since the latter suffers from sampling bias if noise correlations for different location types are not equal, because pairs with high rate map correlations contribute more samples from shared field bins and out of field bins than from one-field bins.

To compare noise correlations between cell-pair types and location types we used an LMM with the equation:

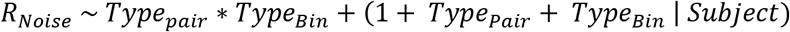

Where *R_noise_* is the average noise correlation for a unit pair at one of the three location types and *Type_bin_* is a categorical variable indicating the location type.

#### Cross-correlogram

Cross-correlations of A1 – V2L unit pairs were computed from the z-scored spiking activity of all spatially stable units, during periods with running speed > 0.06 m/s. Spiking activity was first binned in 10 ms bins. The mean spiking activity observed in the spatial bin occupied by the rat at each instant was then subtracted from the spiking activity in order to prevent the relative spatial positions of firing fields of two units from biasing the shape of the cross-correlogram. In doing so, the cross-correlation only considered activity fluctuations with respect to the mean activity observed at each location. Histograms were made of the distribution of cross-correlogram peak lags for each animal individually, which were then averaged to produce the histogram shown in fig. 3G.

#### Information theoretic analysis

For the information theory-based analyses, linear spatial bins of ∼16.4 cm were used, such that the linearized track was divided into 21 bins. As predictors we used running speed, head direction (θ_head_) and head direction change (Δθ_head_). Running speed and θ_head_ were calculated as described above, and Δθ_head_ was calculated as the degrees per second change in θ_head_. The predictors were binned in 21 equipopulated bins to match the number of location bins. Spike counts were binned in 300 ms time bins, and instances with running speed < 0.1 m/s were excluded. Discrete mutual information (MI) and discrete conditional mutual information (cMI) were computed using the Java Information Dynamics Toolkit (JIDT)(Lizier, 2014). Bias due to finite sample sizes was corrected by generating for every computation of MI and cMI a population of 500 surrogates. When creating the surrogates, only spike count vectors were shuffled, such that the relationship between the target and conditional predictor (in the case of cMI) was preserved. The MI and cMI values reported are the differences between the observed values and the average of the surrogate populations. Differences in MI between A1 and V2L were statistically assessed using Mann-Whitney’s U test. The p-value of cMI values of individual units was computed as the fraction of the surrogate dataset which had information values higher than the observed one. To correct for multiple comparisons, a Bonferroni correction was applied across all neurons and tested at the 0.05 significance level. Reported confidence bounds correspond to 95% bootstrap confidence intervals, computed with the Seaborn Python library (v0.9.0, 2018).

#### Encoding: predicting spike trains from behavioral variables

First, the position of the rat was taken as the two-dimensional variable with x- and y-coordinates with a resolution of 0.205 cm/pixel recorded by the system without further binning. Encoding was performed with a random forest encoder using 100 trees and 5-fold cross-validation with randomized folds (Benjamin et al., 2018). Encoding over time was performed using continuous folds to preserve the order in time. Encoding quality was measured with the Poisson pseudo-R^2^ score and averaged over folds. Statistical comparisons of encoding quality for individual predictors, and comparisons of improvement in encoding quality above all other predictors, were made using Wilcoxon’s signed-rank test and used the average of the A1 and V2L (improvement in) encoding quality for each predictor. Reported confidence bounds correspond to 95% bootstrap confidence intervals, computed with the Seaborn Python library (v0.9.0, 2018).

#### Decoding of animal position

The position of the rat was decoded from the neuronal data recorded from A1 or V2L if a session included at least 16 neurons from that area showing a rate map peak > 2Hz; only those neurons were included for each session. This number was determined to provide a balance between decoding quality and number of included sessions. Spikes were binned in 400 ms bins, and the true position (i.e. the actual position of the rat on the track) at every timeframe was assigned to a spatial bin on the linearized track (total of 35 bins, bin size ∼9.8 cm). When the linear position changed spatial bins within a temporal bin, the position was assigned to the spatial bin which occurred most often within the temporal bin. Running speed was linearly interpolated at the centers of the temporal bins. Samples with speed < 0.1 m/s and with spike count < 5 were excluded. A Bayesian classifier was employed to predict the spatial bin occupied by the rat on the basis of the temporally binned neural data (Davidson et al., 2009). A 5-fold cross-validation routine with shuffling was used, with identical shuffling (i.e. similar sized training set for each fold) across the two areas for a given session.

Decoding errors were used as a main metric for decoding performance and computed as the Euclidian distance between the centers of the true and the decoded spatial bin in 2D space. Pearson correlations of instantaneous decoding errors between the two cortical areas in time were calculated to assess whether A1 and V2L encoded the same position. This was performed separately for the error in the x-direction and that in the y-direction in 2D space, to preserve the directionality of the error in addition to its magnitude.

Instantaneous error correlations resulting from these computations were compared with error correlations computed following shuffling of the errors within the same spatial bin and running speed range (Saleem et al., 2018). Running speed bins were defined per session by taking the full range of speeds and subdividing it into 5 equipopulated bins. Significance of differences in error correlations before and after shuffling were tested using Wilcoxon’s signed-rank tests. Joint error density maps for A1 and V2L were computed for the error correlations of recorded and shuffled data and for the x- and y-direction separately. Joint error density maps were averaged across all included sessions and smoothed with a Gaussian filter with a standard deviation of 4 spatial bins. The relative probability of observing an error of a particular size and direction in the recorded versus shuffled data was calculated by taking the difference between the actual and shuffled joint density maps and dividing by the shuffled map.

A bootstrapping procedure was performed for testing how the decoding performance depended on the size of the population. First, for each included session, 50 unique, random groups of units were selected for each ensemble size (ranging from 5 to the maximum number of units in each session minus one). Then, the decoding analysis was performed for each group before averaging decoding performance across the groups. For the largest ensembles, with one fewer unit than the session total, it was not possible to create 50 unique groups. For these ensembles some groups were included twice. Cross-validation was performed for every group by splitting the data into a training set (80%) and test set (20%), with identical shuffling across all groups of a session.

To exclude the possibility that the observed correlations in instantaneous decoding errors are a result of decoding artifacts in sessions with poor decoding, Pearson’s correlation was computed for the average decoding errors and the correlation in instantaneous decoding errors across sessions, for A1 and V2L average errors separately and for instantaneous errors in the X- and Y-directions separately.

### Seaborn Python reference (no journal article associated)

v.0.9.0 DOI 10.5281/zenodo.1313201, 2018

## Acknowledgements

This study was financially supported by Netherlands Organization for Scientific Research VENI Grant 863.11.010 to C.S.L. and by the European Union’s Horizon 2020 Framework Program for Research and Innovation under the Specific Grant Agreement No. 945539 (Human Brain Project SGA3). We thank Kenneth D. Harris and A. David Redish for the availability of unit isolation software KlustaKwik and MClust, respectively. We also gratefully acknowledge the use of a Seaborn Python library in our study (v0.9.0, DOI 10.5281/zenodo.1313201, 2018). The work of the Technology Center at the University of Amsterdam for building the recording setup and the tetrode microdrives is highly appreciated.

---

**Supplementary figure S1:**
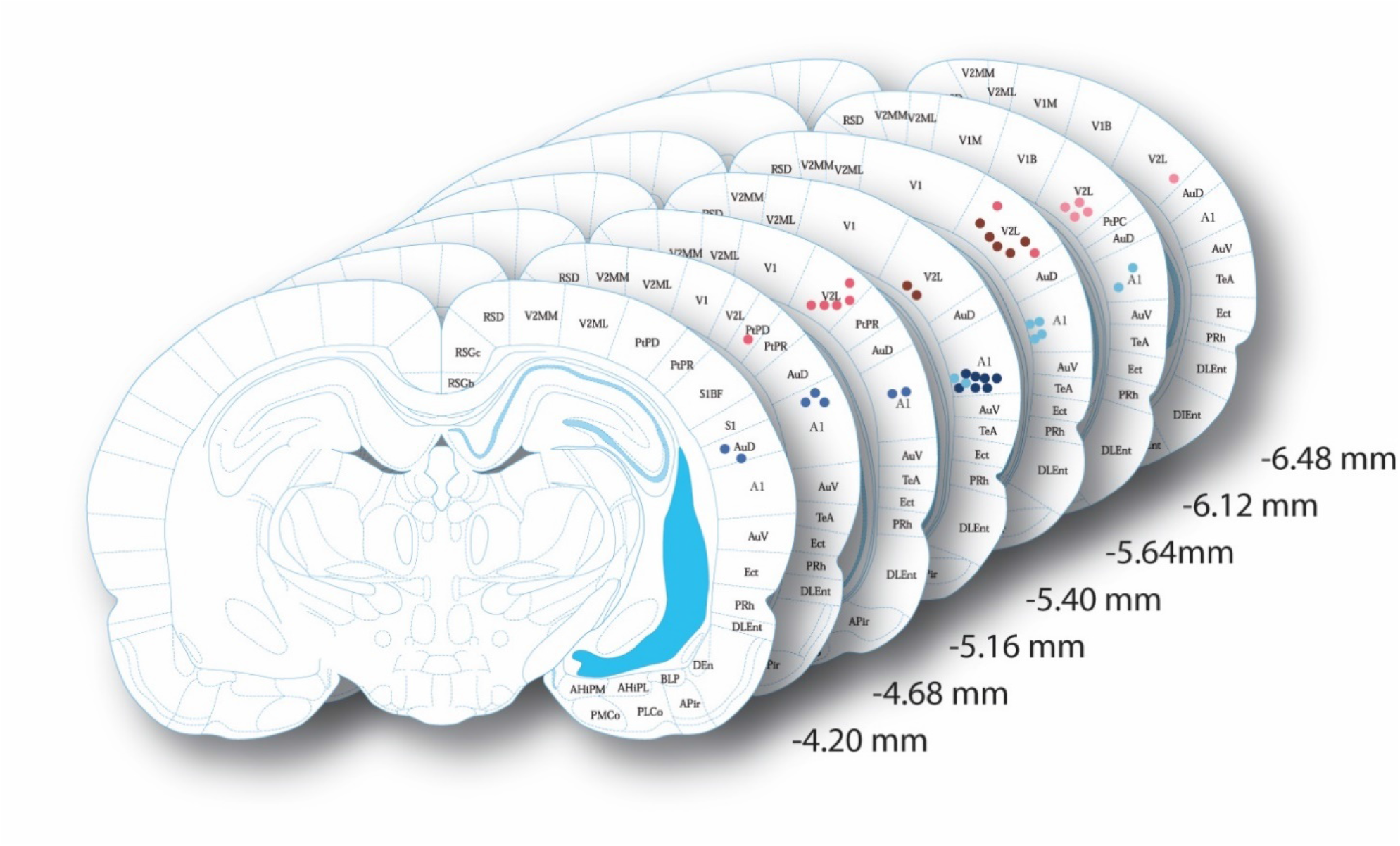
Histology. Dots indicate locations of tetrode endpoints. Red dots represent endpoints of tetrodes targeted at V2L, blue dots endpoints of tetrodes targeted at A1. Three different shades were used to represent data from the three subjects. Light blue and light red correspond to the first subject, medium blue and medium red to the second subject and dark red and dark blue to the third subject. Numbers indicate distance from Bregma. Plates adapted from Paxinos & Watson (2007).

**Supplementary figure S2:**
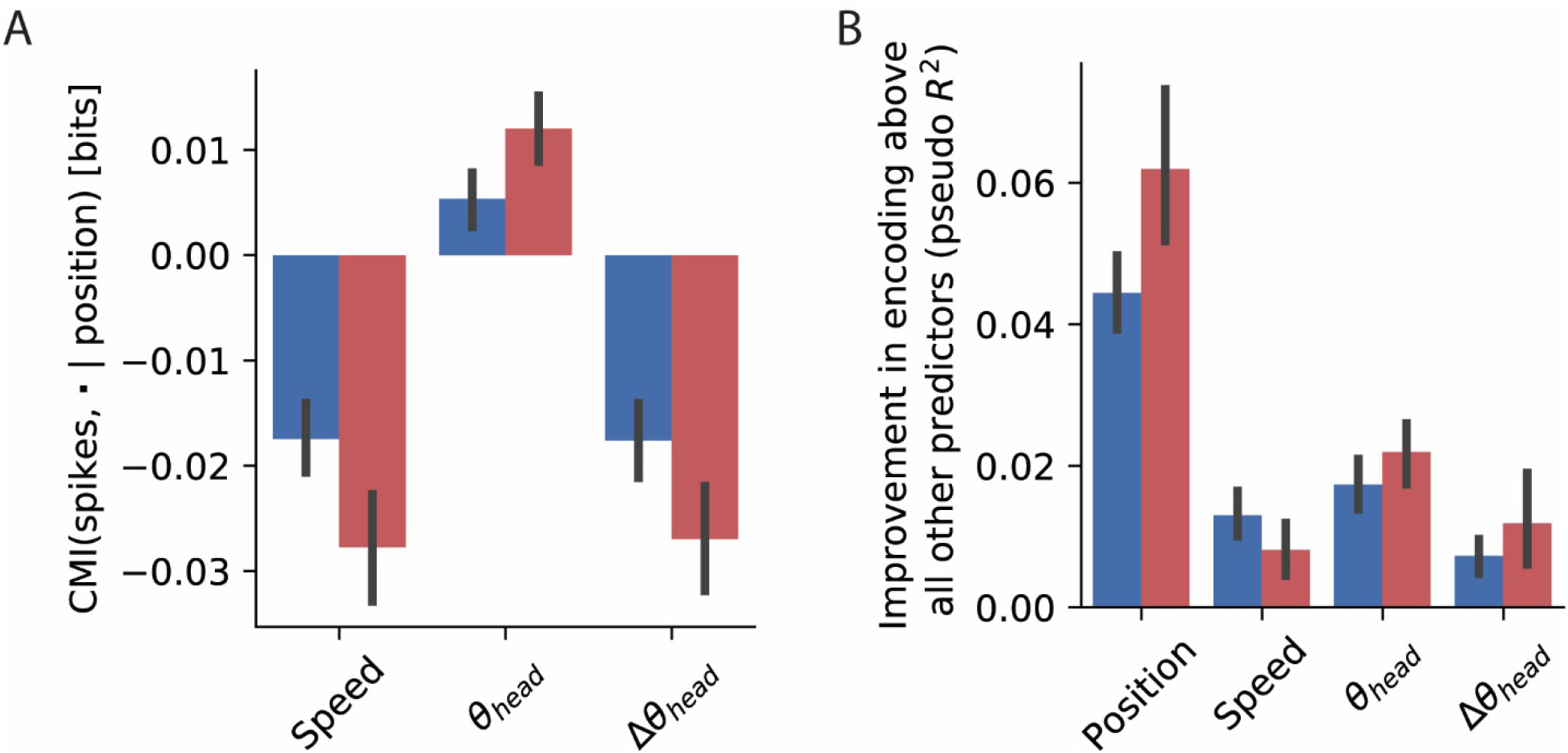
Encoding without stimulus presentation period. **(A)** Debiased, conditional mutual information (cMI) between spiking activity and behavioral factors speed, head direction and changes in head direction, conditional on position (red bars: V2L; blue bars: A1). For this analysis, data from the period around stimulus presentation ± 1 s were ignored. Error bars represent 95% bootstrapped confidence bounds. **(B)** Mean improvement in encoding quality of the random forest encoder across all single units following the addition of the indicated behavioral factor to a model already containing all other factors. For this analysis, data from the period around stimulus presentation ± 1 s were ignored. Error bars are 95% confidence bounds.

**Supplementary figure S3:**
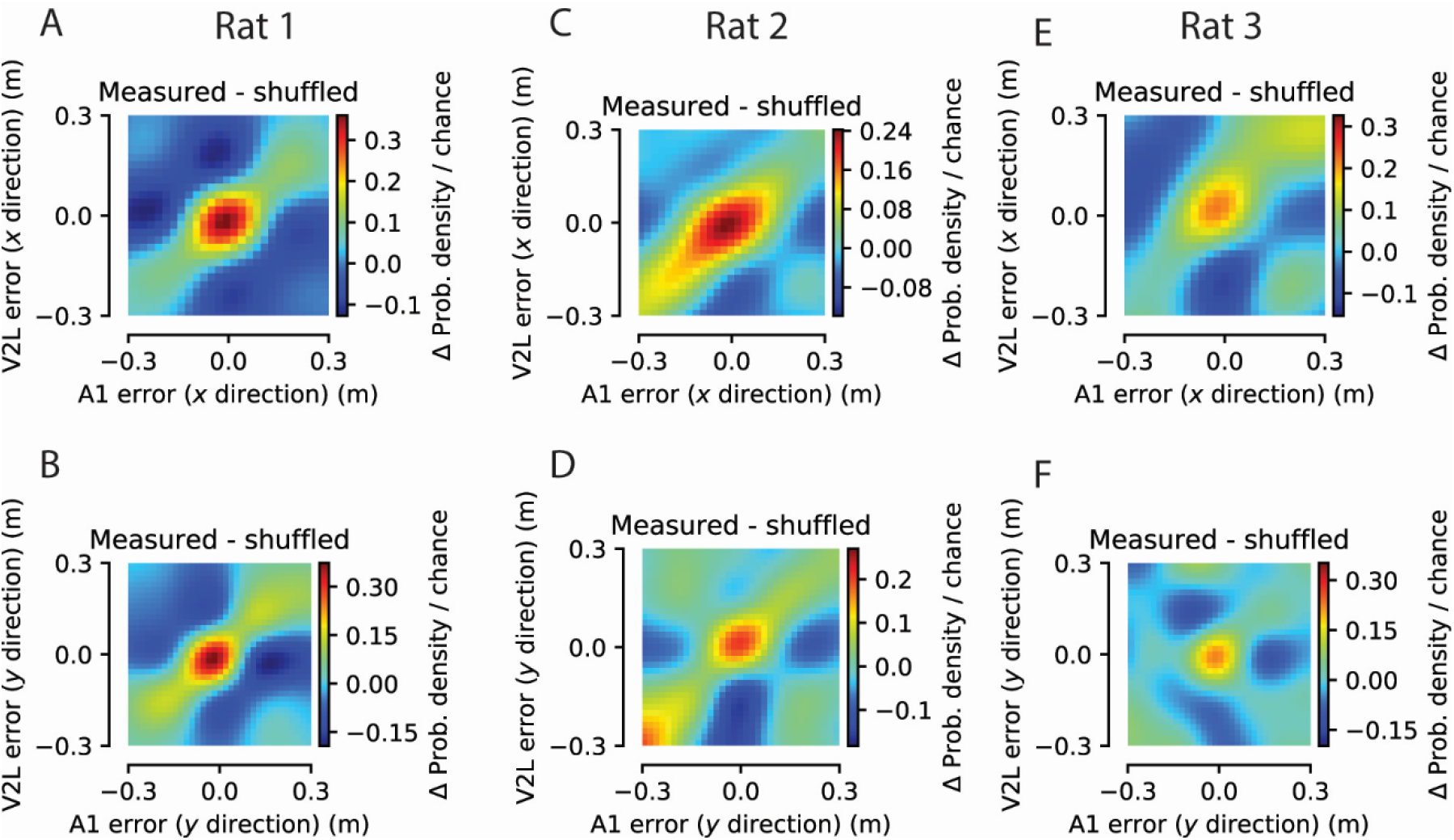
Correlations in instantaneous decoding error per subject. The size and direction of instantaneous decoding errors were correlated between A1 and V2L for individual subjects. **(A)** Color coded is the frequency of observing an instantaneous decoding error of a particular size and direction in the X-dimension (in A1, abscissa, vs. V2L, ordinate) relative to the frequency of instantaneous errors of a particular size and direction following shuffling of the errors within the same spatial bin and speed range. Data from the first subject. **(B)** as (A) but for the Y-dimension. **(C)** and **(D)**, as (A) and (B) but for the second subject. **(E)** and **(F)** as (A) and (B) but for the third subject.

**Supplementary figure S4:**
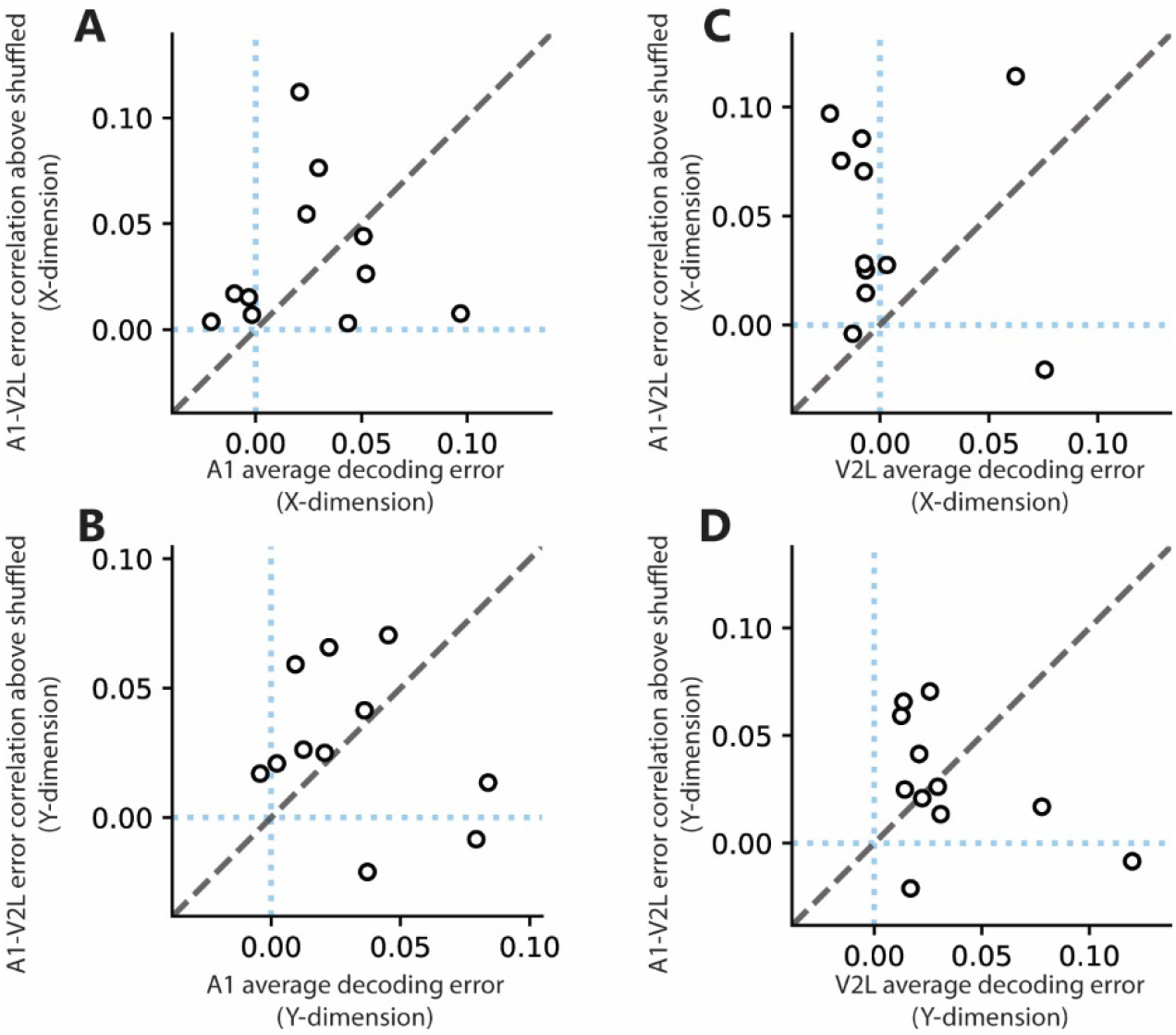
Average decoding error and instantaneous error correlations between areas are uncorrelated. **(A)** Using Pearson’s correlation, we found no linear relationship between the average decoding error in the X-dimension in A1 and the average correlation in instantaneous decoding error in the X-dimension between A1 and V2L. Each dot represents a session included in the decoding analysis for both sessions. **(B)** As (A) but for the Y-dimension. **(C)** As (A) but for the average decoding error in V2L. **(D)** as (C) but for the Y-dimension.

